# Not just mutations: Inbreeding depression persists without genetic variation

**DOI:** 10.1101/2025.02.25.640231

**Authors:** Nicolás Bonel, Christoph Grunau, Patrice David

## Abstract

Inbreeding depression (ID), the decline in fitness upon inbreeding, is thought to result from a decrease in genetic heterozygosity enhancing phenotypic effects of recessive deleterious mutations. However, emerging evidence suggests that mutations may not explain ID completely. In this study, we test whether ID can emerge even in contexts where genetic heterozygosity does not vary. To that end, highly inbred lines (*F*=0.99999997) of the freshwater snail *Physa acuta* were used to produce individuals with varying levels of parental relatedness (self-fertilization, sibling crosses, and cousin crosses), though with identical genomic heterozygosity. Several fitness traits declined significantly with increasing parental relatedness, a pattern characteristic of ID, and quantitatively representing a non-negligible fraction of the ID usually observed in natural, genetically diverse populations of *Physa acuta*. Individual-based simulations showed that mutation rates compatible with values of ID found in natural populations are way too low to generate as much ID as observed in our experiment. These findings are consistent with the hypothesis that epigenetic changes, in addition to mutations, could contribute to a rapid regeneration of ID and explain the persistence of detectable ID in sets of genetically identical individuals.

## Introduction

Inbreeding depression (ID) is the loss in fitness that occurs in offspring of consanguineous matings and is one of the most general observations in evolutionary biology (Charlesworth and Charlesworth, 1999; Charlesworth and Willis, 2009).

Understanding the mechanisms of ID is important because it strongly constrains the evolution of mating systems (self-fertilization rates, self-incompatibility) and is a major concern in plant breeding and animal conservation (Charlesworth and Charlesworth, 1998). ID is classically explained by the presence of recessive deleterious mutations in the genome, whose effects are exposed by homozygosity (Charlesworth and Willis, 2009). Most of the mutations that occur spontaneously at every generation are indeed deleterious and partly recessive (Agrawal and Whitlock, 2011; Charlesworth and Willis, 2009; Huber et al., 2018; Simmons and Crow, 1977). In large diploid organisms, however, genomic mutation rates are typically small, implying that the standing inbreeding load is accumulated through many generations until a steady state of mutation, selection, and drift is reached. This view inspired the idea that a substantial fraction of standing ID could be removed in conditions that drastically reduce heterozygosity by increasing the elimination or fixation rates of alleles. The so-called genetic purging is expected to occur by selection when unusual amounts of systematic inbreeding (such as high selfing rates) are applied to large populations, exposing deleterious alleles to selection (Glémin, 2003; Lande and Schemske, 1985). ID can also be lost by drift when population size is severely reduced (Glémin, 2003; Kirkpatrick and Jarne, 2000), as alleles (deleterious or not) get fixed and heterozygosity is lost.

Inbreeding depression is, however, surprisingly difficult to eliminate completely (Waller, 2021). First, while some evidence of genetic purging has come from comparative studies in natural populations as well as a few experiments (Barrett and Charlesworth, 1991; Carr and Dudash, 1997; Chelo et al., 2014; Noël et al., 2016a), some residual ID frequently persists even in populations subject to recurrent inbreeding (Byers and Waller, 1999; Carr and Dudash, 1997; Chelo et al., 2014; Crnokrak and Barrett, 2002) Second, ID is also not systematically eliminated by drift, even within lines with extremely reduced effective size and highly homozygous, such as those propagated by continuous sib-mating or continuous self-fertilization (e.g., Han et al., 2021). A classical argument explaining the difficulty of purging refers to the genetic architecture of the mutation load. If ID relies on the joint effect of many loci with very small effects, allele frequency changes under selection are expected to be very slow (Charlesworth and Charlesworth, 1990; Lande et al., 1994). This argument, however, does not apply to fixation by drift. Purging and fixation may also be impeded by linkage and selective interference between loci (Lande et al., 1994). Indeed, linkage disequilibrium among deleterious mutations may preserve heterozygosity (and ID) in some regions of the genome, when two deleterious alleles with strong effects (such as lethals) are in repulsion, or when stretches of weak-effect deleterious mutations collectively form pseudo-overdominant blocks (Waller, 2021). Thus, linkage can delay the loss of heterozygosity in small populations and inbred lines. However, this loss eventually takes place given sufficient time, as recombination ends up breaking the associations that prevent homozygosity. Once homozygosity is attained in a genomic region, the sole mechanism that may reintroduce heterozygosity, and regenerate ID, is the occurrence of spontaneous mutations. Mutations, however, are rare and, within a selfing or sib-mating line, they are quickly lost or fixed, so they are not expected to yield substantial levels of ID. Overall, an extended period of very low effective population sizes, such as in an anciently established inbred line, makes it unlikely to observe significant levels of ID. In other words, aside from a few recent mutations, two individuals from the same inbred line are essentially genetic clones. As a result, crossing them is genetically equivalent to self-fertilization, and the fitness of offspring from crossbreeding and selfing should not differ.

Recent studies, however, suggest another process that could regenerate inbreeding depression in genetically uniform lines faster than spontaneous mutation: epigenetic change (reviewed by Cheptou, 2024). Support for a role of epigenetics in ID comes from several lines of evidence (Biémont, 2010; Bossdorf et al., 2008; Cheptou and Donohue, 2013; Vergeer et al., 2012). First, ID can be greatly reduced by applying a demethylating enzyme (Vergeer et al., 2012). Second, in maize, the continued decline in performance within selfing lines from generations 7 to 11 is accompanied by a global increase in DNA methylation while residual heterozygosity is extremely low (Han et al., 2021). This suggests that epigenetic change can co-occur with sustained phenotypic deterioration even when genetic homozygosity itself does not increase anymore.

Similarly, in *Arabidopsis,* heterosis (the reciprocal of ID) is observed when crossing isogenic epi-RILs (i.e., selfing lines differing only in methylation patterns; Dapp et al., 2015).

Epigenetic modifications can be independent of genetic variation (“pure” or autonomous epigenetic variants, *sensu* Richards, 2006) thus being a possible source of heritable phenotypic variation in the absence of DNA sequence change (Johannes et al., 2009). A growing body of evidence suggests that incomplete reprogramming and erasure of epigenetic marks during gametogenesis and zygote development could promote stable transgenerational transmission of some epigenetic variants (Preite et al., 2018). Epimutations are generated at higher rates than genetic mutations and tend to revert to the wild-type in a few generations (Van Der Graaf et al., 2015). A fraction of the standing ID could therefore reflect fast-generated epigenetic changes transmitted over one or a few generations. Phenotypic consequences of autonomous epigenetic inheritance have been classically studied using systems with a known lack of genetic variation (e.g., Bossdorf et al., 2008). In a similar way, if autonomous epigenetic changes play a role in ID, differences in offspring fitness should persist among different types of crosses (self-fertilized, sib-mating, cousin-mating) even when all parents used are genetically identical (i.e., clonemates). Yet, this approach has not been undertaken yet.

Here we tested whether significant ID can be detected even when differences in genomic heterozygosity are extremely low. To that end, we created “outbred clones” by intercrossing inbred lines. We produced two highly inbred lines of the hermaphrodite freshwater snail *Physa acuta* (a naturally outcrossing species) that have undergone more than 28 generations of enforced self-fertilization, ensuring minimal residual heterozygosity. We chose this species because it is easy to raise in the laboratory, ID has been repeatedly observed in its natural (outbred) populations, and it can be forced to self-fertilize (the most efficient way to eliminate heterozygosity). Individuals within each line are genetically homozygous and identical, except for neo-mutations in the last few generations. We crossed individuals from two different inbred lines and obtained a set of offspring that were heterozygous and identical to one another (“outbred clone”).

We then performed different types of matings with individuals from this outbred clone: self-fertilization, sibling pair-crossing, and cousin pair-crossings. These treatments yielded second-generation offspring with the same expected genomic heterozygosity (ignoring recent mutations), but that differed in pedigree inbreeding, i.e., the number of generations tracing back to the common ancestor of the two uniting gametes. We chose to start with outbred parents (though genetically belonging to the same “clone”) rather than directly from inbred ones, because highly inbred individuals are rather weak, as expected, a condition we deemed “unnatural”. In contrast, the results obtained with normally heterozygous, outbred individuals, are representative of processes that could contribute to ID in natural contexts.

To evaluate whether parental relatedness (mother and father being either the same individual, or siblings, or cousins) affected progeny fitness, we measured juvenile survival, adult body size, probability of laying eggs (in isolation) at the age of 50 days, and female fecundity after insemination. Classical genetic theory predicts the same average performances in all three types of parental relatedness as offspring all have the same genomic heterozygosity.

## Results

Figure 1 recapitulates the pedigree of the parents (F_2_) we used to produce F_3_ snails by either self-fertilizaton, sib-mating or cousin-mating. Table 1 and Figure 2 summarize our main results, while linear models are reported in detail in Tables S1-S3.

**Figure 1.**
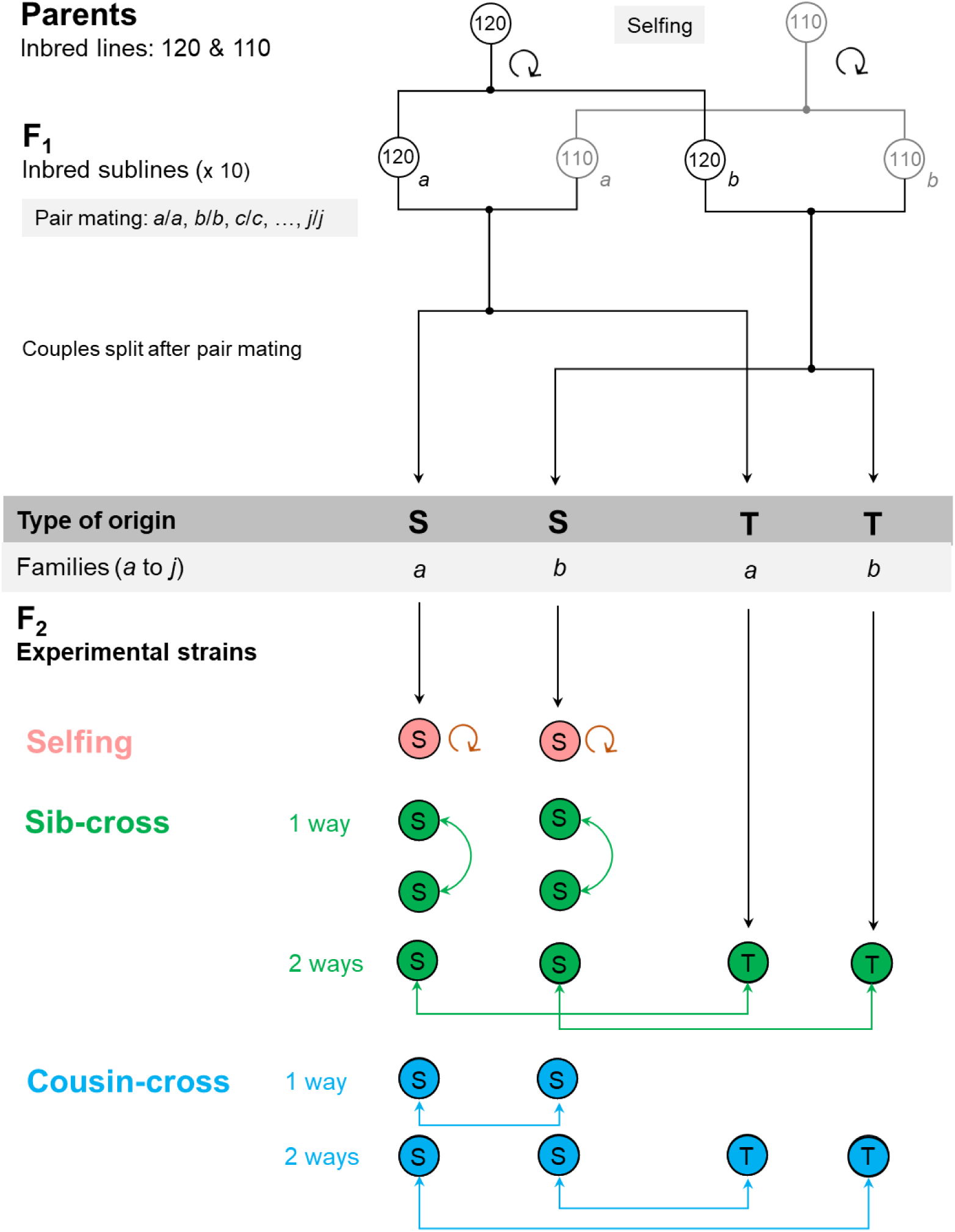
Pedigree of the F_2_ snails used in this study, and how they were crossed to produce the F_3_ on which fitness traits were measured. Parents were two individual snails extracted from independent inbred lines (named 120 and 110) derived from wild-caught ancestors throughr 28 or 29 successive generations of self-fertilization in the laboratory, respectively. F_1_ individuals (inbred sublines) resulted from self-fertilization of parents, and were also homozygous. The F_1_ pair crossing yielded a set of individuals that constituted the experimental strains (F_2_ snails), which are all heterozygous and clonal to one another but with different type of origins, S and T. F_2_ snails from the S type had their mothers from the 120 line and their fathers from the 110 line, and *vice versa* for those from type T. The experimental strains were constituted by ten families (*a* to *j*) per type of origin.

**Figure 2.**
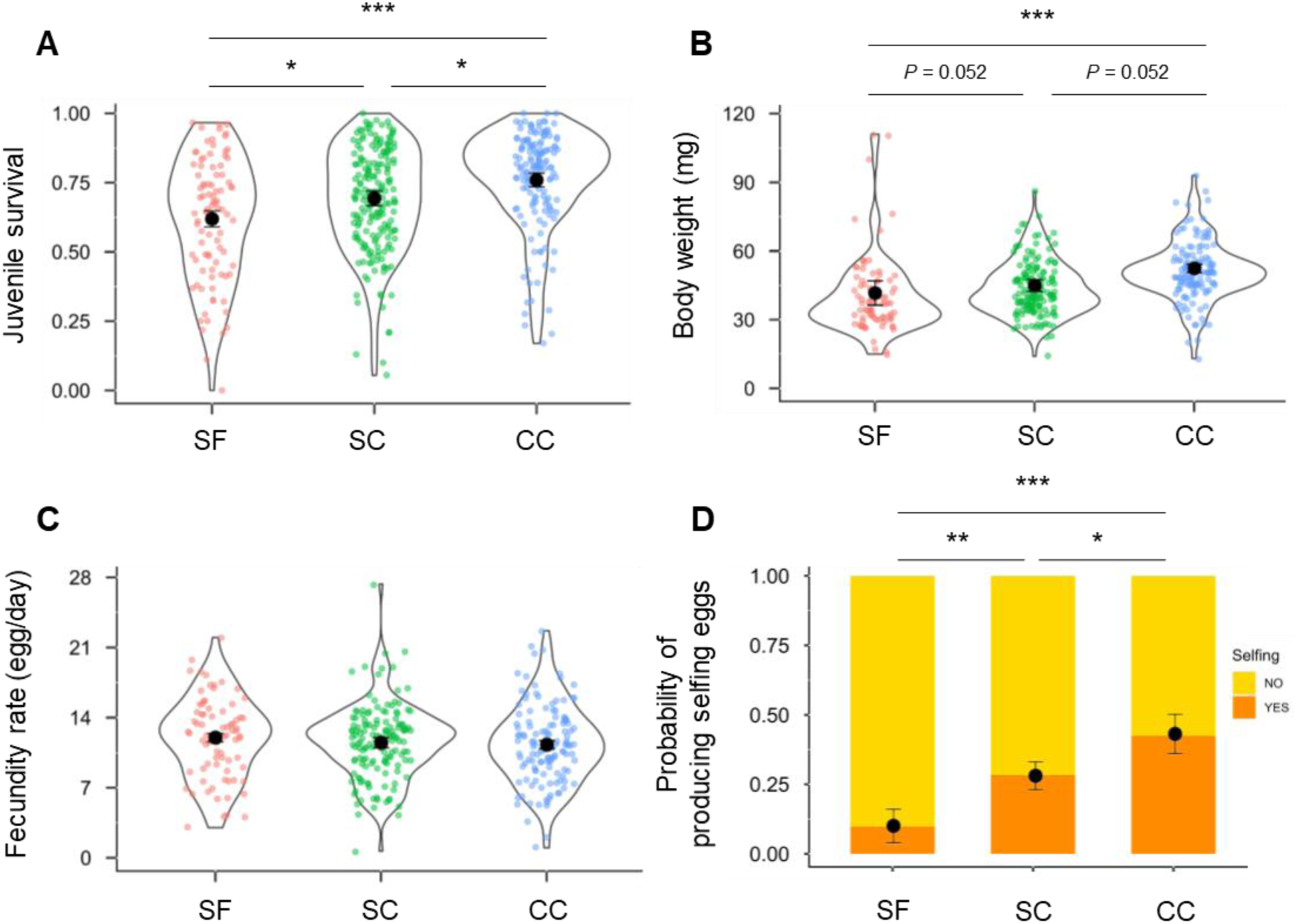
Fitness traits measured on F_3_ individuals of the hermaphrodite snail *Physa acuta*. **A.** Juvenile survival (proportion of eggs that become juveniles alive after 14 days). **B.** Body weight at 50-53 days (the mean of the two measures, before and after pairing the focal snail with a virgin albino partner). **C.** Female fecundity rate after insemination (number of eggs per day produced by the focal snail). **D.** Probability of having produced self-fertilized eggs at 50 days just before pair mating. Mating treatment effect sizes (difference between two types divided by its SD) were tested using post hoc tests with the Holm-Bonferroni correction for multiple-testing (^∗^*P* < 0.05; ^∗∗^*P* < 0.01; ^∗∗∗^*P* < 0.001). SF, selfing; SC, sib-cross; CC, cousin-cross. Black dots are mean values ± 1SE. We did not separate one-way and two-way crosses in the SC and CC category as differences between them were small and not significant (see supplementary tables S1-S3).

**Table 1.** Summary of means (±SE) and statistical significance of the mating treatment type effects on fitness traits measured on F_3_ individuals of the hermaphrodite snail *Physa acuta*. Number of observations are indicated in parentheses. In this table, as in fig.2, we did not distinguish, within ecah mating treatment, the different possible types of crosses (one-way versus two-way, see fig.1) because they were not statistically different. Detailed results are reported in supplementary tables S1-S3.

| Variable | Mating treatment |  |  | Treatment effect | Treatment comparison | Effect size |
| --- | --- | --- | --- | --- | --- | --- |
|  | Selfing (SF) | Sib-cross (SC) | Cousin-cross (CC) |  |  |  |
| Juvenile survival | 0.62 ± 0.03 (93) | 0.69 ± 0.03 (193) | 0.76 ± 0.02 (153) | $X^2_2 = 15.40$; $P < 0.001$ | SF vs. CC | -4.04*** |
|  |  |  |  |  | SF vs. SC | -2.74* |
|  |  |  |  |  | SC vs. CC | -2.33* |
| Body weight | 41.6 ± 5.3 (82) | 44.9 ± 2.6 (152) | 52.3 ± 1.8 (134) | $X^2_2 = 10.06$; $P = 0.007$ | SF vs. CC | -2.98** |
|  |  |  |  |  | SF vs. SC | -1.94 . |
|  |  |  |  |  | SC vs. CC | -2.23 . |
| Fecundity rate | 12.0 ± 0.4 (79) | 11.5 ± 0.5 (146) | 11.3 ± 0.4 (127) | $X^2_2 = 1.53$; $P = 0.466$ | — | — |
|  |  |  |  |  | — | — |
|  |  |  |  |  | — | — |
| Probability of producing selfed eggs at the age of 50 days | 0.10 ± 0.06 (82) | 0.28 ± 0.05 (152) | 0.43 ± 0.07 (134) | $X^2_2 = 19.69$; $P < 0.001$ | SF vs. CC | -4.25*** |
|  |  |  |  |  | SF vs. SC | -3.09** |
|  |  |  |  |  | SC vs. CC | -2.0 * |

Treatment means ( ± SE, with ) are reported in natural scale for the four traits (proportion of eggs surviving, milligrams, eggs laid per day, and proportion of laying individuals respectively); however statistical models are generalized linear mixed models on the appropriate scale (Binomial for juvenile survival and probability of producing selfed eggs, with logit link; Gaussian for body weight and fecundity, identity link), and include appropriate random effects to represent the structure of our design (not shown here, see methods and detailed results in Supplementary materials and tables S1-S3). Mating treatment effect sizes (estimated mean difference on transformed scale divided by its SD) were tested using post hoc tests with the Holm-Bonferroni correction for multiple testing. \* $P < 0.05$ ; \*\* $P < 0.01$ ; \*\*\* $P < 0.001$ after correction.

Juvenile survival significantly differed across treatments. The highest survival was in the offspring of cousin-cross (average 0.76) whereas survival in the selfing and sib-cross treatments was reduced by 18% and 9%, respectively, compared to cousin-cross (Table 1, Fig. 2A). Body weight showed a similar pattern, the cousin-cross treatment had the highest mean, with a reduction of 20% and 14% in the self-fertilization and sib-cross treatment, respectively (Table 1, Fig. 2B). Despite this difference in body weight, there was no significant effect of parental relatedness on female fecundity at 53 days, after insemination (Table 1, Fig. 2C). In contrast, we found a strong treatment effect on the probability of having produced eggs by self-fertilization at the age of 50 days. This probability was the highest in cousin-cross treatment and it decreased by 77% and 35% in the selfing and sib-cross treatments, respectively (Table 1, Fig. 2D).

We genotyped F_2_ individuals, which represent inter-line crosses in our design (their offspring are F_3_), using microsatellites fixed for alternative alleles in the two inbred lines. As expected, almost all F_2_ were fully heterozygous, with rare exceptions where one or two loci appeared homozygous in a few individuals, likely due to genotyping errors (Fig. 3). In the F_3_ generation, we found no significant differences in heterozygosity among mating treatments (*X^2^_2_* = 0.475, *P* = 0.789, binomial model on the proportion of heterozygous loci). The multilocus heterozygosity distribution within each treatment was centered around 0.5, consistent with expectations for F_2_×F_2_ progeny: selfing (*H_O_* = 0.51±0.05), sib-cross (*H_O_* = 0.49±0.16), and cousin-cross (*H_O_* = 0.52±0.16; Fig. 3).

**Figure 3.**
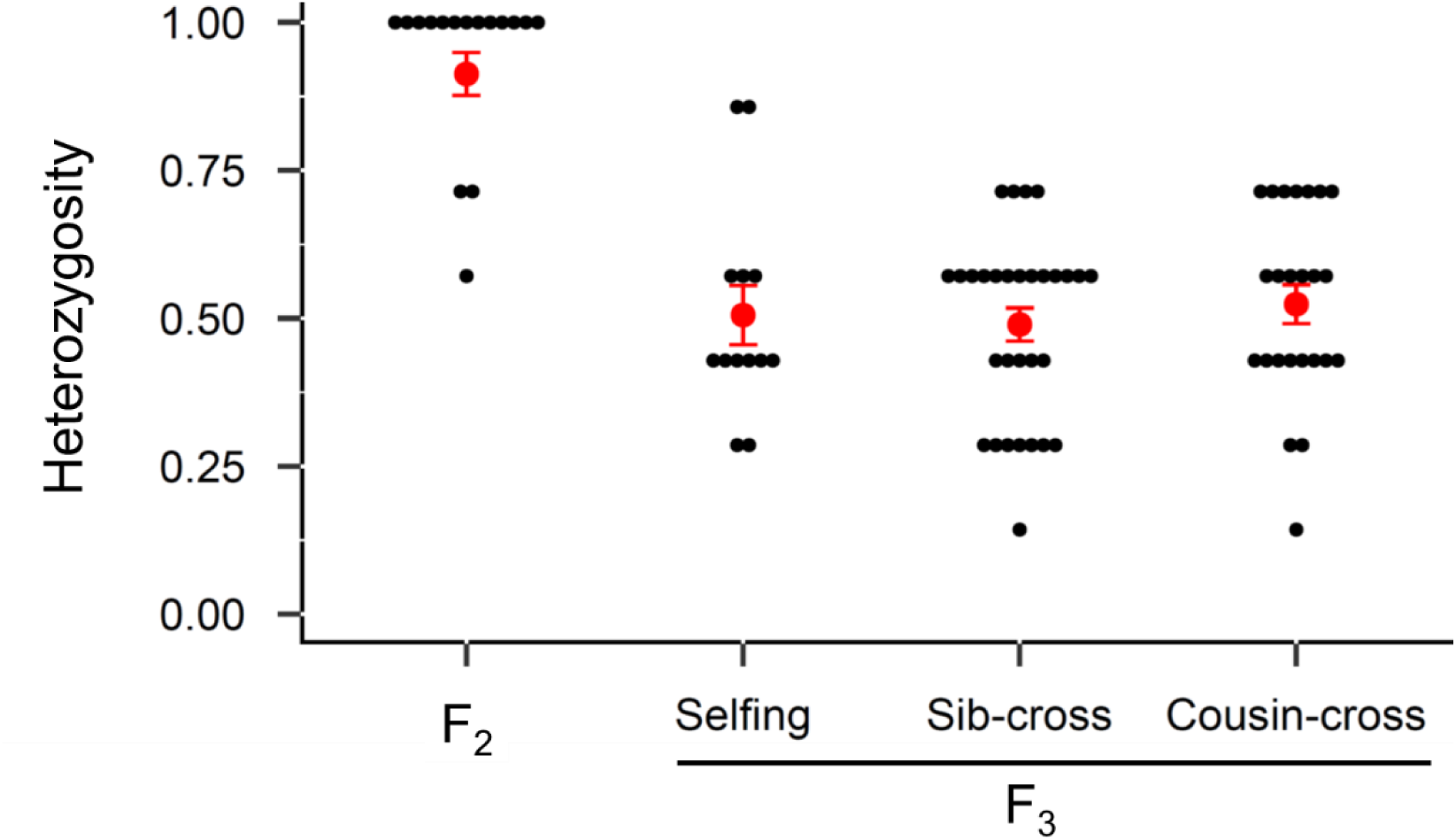
Distributions of multilocus heterozygosity for a subset of F_2_ and F_3_ individuals of different categories using 7 microsatellite loci (AF108762, AF108764, Pac1, Pac2, Pasu1-2, Pasu1-9, Pasu1-11) chosen because preliminary data indicated that the inbred lines 110 and 120 were fixed for different alleles. The microsatellites were grouped into two multiplexes. Overall, F_2_ individuals were fully heterozygous (with a few exceptions) whereas in the F_3_ heterozygosity was spread around 0.5 within each mating treatment. Red dots are mean values ± 1SE.

We conducted simulations replicating our experimental design to determine whether the observed differences between treatments could result from incomplete fixation, linkage, or spontaneous mutation. The mutational parameters were selected to align with previous estimates of ID in natural, outbred populations of *Physa acuta*(Escobar et al., 2008). The simulations showed that offspring fitness from full-sib and cousin crosses was only slightly higher than that from the selfing treatment, with differences of 0.8% and 2.0%, respectively—well below the observed differences in juvenile survival, body weight, and the probability to lay eggs in isolation (Fig. 4). We tested various parameter sets compatible with naturally observed levels of ID, adjusting linkage, contributions of lethal mutations, and selection or dominance coefficients, but these had minimal effects on the results. The largest predicted difference in fitness between cousin-cross, sib-cross, and self-fertilization occurred when the source of ID was driven quasi-exclusively by mutations to semi-lethals (*d_1_*=0.49, *d2*=0.01) and when these mutations were nearly fully lethal (*s_1_*=0.99). However, even under these extreme conditions, average fitness differences never exceeded 5% (see Fig. S7). While individual replicate runs varied considerably within each parameter set, the mean predictions consistently fell well below the observed ratios in all situations. Simulations could match the experimental results only when the mutation rate was artificially increased by approximately 10-to 12 fold beyond the levels of ID estimated for natural populations of *P. acuta* (Fig. S8).

**Figure 4.**
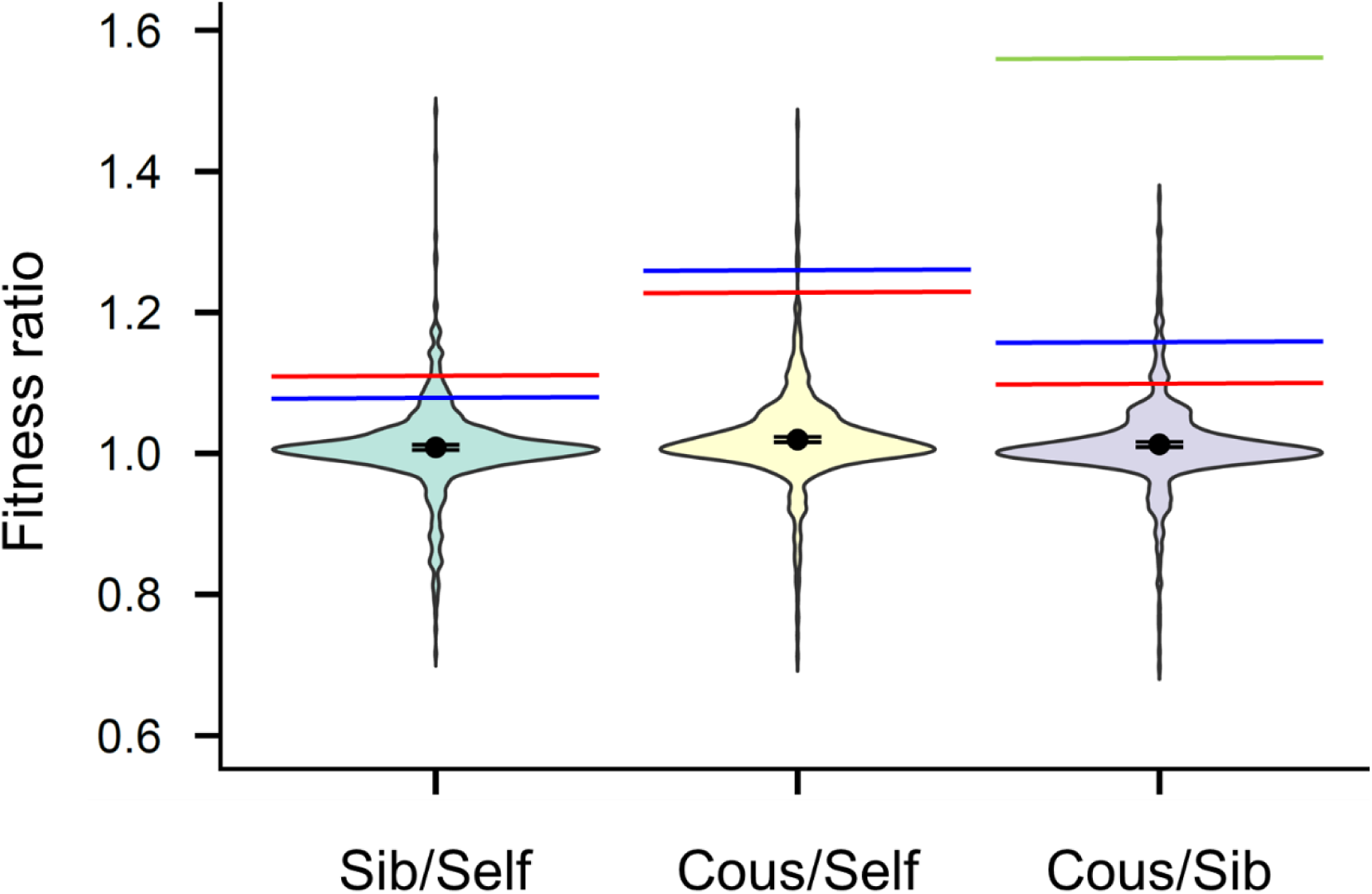
Violin plots of the distribution of fitness ratios from 1,000 simulations of our experiment, using the reference parameter set, compatible with available data on ID in *Physa acuta* (see Methods and *Simulations* in Supporting Information). Parameters include the length of the genome (*L* = 10 Morgans), the dominance coefficient and effect size of semi-lethals (*h_1_* = 0.02, *s_1_* = 0.9), and those of small-effect mutations (*h_2_* = 0.2, *s_2_* = 0.05), with both mutation types contributing to a total inbreeding depression of *d=0*.5 in natural populations (*d_1_* = 0.3 for semi-lethals and *d_2_* = 0.2 for small-effect mutations). Black dots represent the mean fitness ratio, with 95% confidence intervals (CI) shown. The observed ratios for juvenile survival (red line), body mass (blue line), and the proportion of fertile individuals at 50 days (green line) were for Sib/Self: 1.11, 1.08, and 2.80 (not shown, out of range); for Cous/Self: 1.23, 1.26, and 4.30 (not shown); and for Cous/Sib: 1.10, 1.16, and 1.54, respectively. Simulations with different parameter sets are in Figure S7 from Supporting Information.

## Discussion

Our study reveals substantial inbreeding depression (ID) in the progeny of experimental lines, characterized by distinct levels of parental relatedness. Three fitness traits—juvenile survival, body mass, and self-fertility—declined as parental relatedness increased (self<sibs<cousins), while female fecundity showed no significant differences. The amount of ID in juvenile survival (0.18) and body mass (0.20) in the selfing treatment, relative to the cousin cross, is lower than typical values reported for outbred and self-fertilized offspring of wild-caught *Physa* acuta, where ID estimates are approximately 0.55 for juvenile survival (Escobar et al., 2008; Janicke et al., 2013) and 0.38 for body mass (Janicke et al., 2013). Comparable data for self-fertility are not available. Despite these lower ID values, the levels observed in our experiment— amounting to one-third to one-half of those in natural outbred populations for key traits—are striking and somewhat paradoxical given the unique design of our study: all three types of offspring have identical genomic homozygosity. During the establishment of the inbred lines, deleterious mutations have been either eliminated or fixed, resulting in identical levels of heterozygosity across all F_3_ categories (selfed, sib-crossed, and cousin-crossed)—all expected to retain half the heterozygosity of the genetically uniform F_2_. Under mutation-based ID theory, we would predict similar average fitness across treatments, yet our results clearly indicate otherwise.

One potential explanation for this result is that some F_2_ or F_3_ individuals may have failed to successfully cross-fertilize, possibly due to a loss of cross-fertilization ability after 28 generations of enforced selfing. However, two key arguments dispel this possibility. First, additional crosses confirmed that these individuals could both fertilize and be fertilized by their partners (see Supporting Information: *Outcrossing ability of inbred lines*). Second, microsatellite genotyping supported the expected patterns, with F_2_ individuals showing full heterozygosity for the diagnostic markers. Occasional apparent homozygosity at one or two loci is likely due to reading errors and does not indicate selfed offspring, which would be fully homozygous like their parents.

Moreover, observed microsatellite heterozygosity did not differ among the three classes of F_3_ offspring, and was consistently at 0.5, as expected. Note that in classical ID experiments, self-fertilized broods typically exhibit half the heterozygosity of outcrossed ones.

In the absence of differences in genomic heterozygosity, what explains the observed ID? A first hypothesis is that the reduced fitness in the selfing treatment may stem from maternal effects (McCarthy and Sih, 2008). However, such maternal effects in this case would not be linked to variation in maternal quality, as mothers were randomly assigned to all treatments. Instead, they might reflect a parental component of ID, where less energy is allocated to self-fertilized eggs compared to outcrossed ones (e.g., Cheptou, 2024). While plausible, this argument fails to account for the fitness differences between the sib-and cousin-cross treatments. Moreover, fitness traits in *Physa acuta*, including juvenile survival, are not known to be strongly influenced by maternal effects. For instance, outcrossed offspring of inbred mothers do not exhibit lower fitness than offspring of outbred mothers (*cf*. Janicke et al., 2013; Noël et al., 2016a), even though the inbred mothers themselves are less fit than outbred ones.

Alternatively, the observed variation in ID among mating treatments could result from the long-term persistence of localized heterozygosity in specific genomic regions within our inbred lines. This may occur due to linkage disequilibrium among deleterious mutations, constituting pseudo-overdominant blocks (Waller, 2021). Such blocks would not be detected by the seven microsatellites used in our study, which represent only a small fraction of the genome. However, our simulations do not support the hypothesis that pseudo-overdominance, driven by tightly linked deleterious alleles, could persist or accumulate over the 25+ generations of extreme inbreeding experienced by our lines.

Specifically, our simulations show that mutations generate negligible fitness differences among offspring from selfing, sib-, and cousin-crosses. Even when we artificially increased linkage by reducing the genome size to the unrealistic value of 1 Morgan (despite *Physa acuta* having 19 pairs of chromosomes), the resulting fitness ratios remained virtually unchanged. Classical models suggest that pseudo-overdominance is most likely to emerge under conditions of inbreeding combined with very small genome sizes (Waller, 2021; Wang and Hill, 1999; Zhao and Charlesworth, 2016). However, our experimental design involves much more extreme and prolonged inbreeding than most empirical studies that have reported persistent heterozygosity or inbreeding depression in inbred populations (Chelo et al., 2019; Latter, 1998; Rumball et al., 1994; Williams et al., 2016). Under such intense inbreeding, residual heterozygosity can only be maintained in chromosomal fragments harboring deleterious mutations that function as balanced lethals (Wang and Hill, 1999). In this scenario, heterozygosity would be expected to differ between selfed and outcrossed F_3_ offspring, as F_2_ individuals could inherit distinct chromosomes from the F_1_ parents. In contrast, no difference in heterozygosity should be observed between sib-crossed and cousin-crossed F_3_ individuals, since all F_1_ parents would share identical heterozygosity at such pseudo-overdominant chromosome (balanced lethality). Taken together, these considerations suggest that linkage among deleterious mutations is unlikely to produce the patterns of heterozygosity—and corresponding fitness variation—observed in our highly inbred lines.

Simulations suggest that fitness contrasts are maximized when inbreeding depression arises from spontaneous semi-lethal mutations. While these mutations may not persist for long, they can occasionally be present in a heterozygous state in the parents at the time of experimentation. It is important to note that lethal mutations are lost at a rate of one-third per generation in selfing lines. Indeed, homozygous lethal individuals are unviable, so that among viable offspring produced by a heterozygote, one-third are homozygous wild-type, and two-thirds heterozygous. As a result, the half-life of lethal mutations in selfing lines is approximately 2-3 generations. Heterozygosity resulting from mutations with milder effects has an even shorter half-life, as homozygous mutants are not completely eliminated from the population. Thus, in long-established selfing lines, the heterozygous load is likely to reflect only a few generations (2-3) of accumulated spontaneous mutations. However, even in the most favorable scenario, which assumes an unusually high proportion of lethal mutations, our findings remain highly unlikely. In our reference scenario, we modeled an average per-haploid genome rate of spontaneous semi-lethal mutations of *U_1_* = 0.028. Although direct estimates of *U_1_* are unavailable for *Physa acuta*, this value is close to those reported for *Drosophila melanogaster*: 0.003 to 0.017 for the large chromosomes 2 and 3, and 0.006 to 0.034 for the full genome (Charlesworth et al., 2004; Fry et al., 1999; Sharp and Agrawal, 2012). Across all simulations, cousin-crossed or sib-crossed offspring consistently showed, on average, a small but positive viability advantage (<5%) over self-fertilized offspring. This aligns with the persistence of spontaneous mutations over two or three generations, as discussed earlier. However, this predicted advantage is far smaller than the observed differences in juvenile survival. Notably, fitness ratios across simulations exhibit significant stochastic variation (Fig. 4), particularly when lethal mutations are frequent. This variability is expected, given the stochastic nature of recent mutations in the lines just a few generations before the experiment. While the observed survival estimates remain improbable, a small fraction of simulations occasionally yield survival ratios that are similar to those observed: 3.8 % for Sibling/Selfing, 0.8% for Cousin/Selfing, and 6% for Cousin/Sibling (Fig. 4). To definitively rule out a purely mutational origin for these survival differences, replication with independent line pairs would be necessary.

Body size and the proportion of self-fertile individuals at the age of 50 days also showed clear signs of ID, with ratios even higher than those observed for juvenile survival. These traits expressed later in life, typically contribute less to overall ID than juvenile survival in wild-caught *Physa acuta* and are less susceptible to purging (Escobar et al., 2008; Janicke et al., 2013; Noël et al., 2016a). This suggests that the genetic load affecting these traits is more strongly influenced by small-effect alleles than by semi-lethal mutations. Consequently, simulating these traits may require parameter adjustments, such as lower per-generation genomic mutation rates and a higher proportion of small-effect mutations. However, such adjustments would inevitably reduce residual heterozygosity within the lines, further decreasing all ratios and pushing simulated values even farther from the observed results.

Taken together, the unexpectedly high ID observed across three independent traits in our experiment is highly improbable under a mutational genetic model. To reconcile our findings with such a model, one would need to assume a strong (> 10-fold) increase in mutation rates in the inbred lines compared to the original populations (see Fig. S8). While there is evidence that mutation rates can increase in individuals in poor condition, as may be the case in our inbred lines, the magnitude required remains implausible. For instance, studies in *Drosophila* have shown that individuals carrying a load of fixed deleterious mutations—those in poor condition—tend to accumulate spontaneous mutations at higher rates than healthier individuals (Agrawal and Wang, 2008; Sharp and Agrawal, 2012). However the observed increases in mutation rates were modest (up to 2-fold) and did not extend to lethal mutations (Sharp and Agrawal, 2012).

Our results therefore suggest either an increase in mutation rates much higher than previous measures, or a source of fast-renewed ID other than mutation. Differences in fitness-related traits could instead arise from the accumulation of deleterious epimutations, such as DNA methylations, small RNAs, and histone modifications. For instance, in maize, inbred lines exhibit a persistent decrease in fitness alongside a consistent rise in genomic methylation over generations of inbreeding (Han et al., 2021), likely due to disruptions in methylation resetting mechanisms. However, this is unlikely in our study since the founders (F_2_) of the experimental lines were all highly outbred individuals, a condition that would typically restore the overall methylation level back to normal (Han et al., 2021). Instead, our findings support the hypothesis that the observed ID may be driven by epigenetic similarity between uniting gametes.

Epigenetic marks can be transmitted across generations with incomplete fidelity, suggesting that this information is stored within gametes through mechanisms that remain poorly understood (but see Cecere, 2021), and references therein for the role of small RNAs in epigenetic inheritance). Two gametes produced by the same individual are expected to be more similar in their epigenetic information than two gametes produced by full-sibs, while the latter are more similar than those of cousins. This gradient of similarity arises from the increasing number of meiotic cycles, along with subsequent zygote development and (incomplete) epigenetic resetting events, separating paternal and maternal gametes from selfing to sib-cross to cousin-cross (Fig. 1).

Although the above hypothesis has not been previously tested, some studies suggest that increased fitness (heterosis) can result from epigenetic dissimilarity between parents, as observed in the crossing of isogenic epi-RILs in *Arabidopsis* (Dapp et al., 2015). In *Physa acuta*, as well as in other mollusks, parental environmental history can influence offspring’s traits through transgenerational plasticity (e.g., Fallet et al., 2020; Luquet and Tariel, 2016; Tariel et al., 2020). In *Biomphalaria glabrata*, a freshwater pulmonate gastropod closely related to *Physa*, DNA methyltransferase inhibitors have been shown to disrupt methylation patterns, leading to changes in growth and gene expression that persist in the untreated progeny of exposed individuals (Luviano et al., 2021). Such environmental effects may contribute to epigenetic differences in gametes produced by different individuals. More broadly, considering that epimutations are generated at higher rates than genetic mutations (Van Der Graaf et al., 2015), it is plausible that a fraction of the standing ID originates from rapidly generated epigenetic changes. These changes, transmitted over one or a few generations, could lead to a decline in performance as parental relatedness increases. While this specific form of ID has not been explicitly modeled, several theoretical frameworks have explored the dynamics of partially recessive deleterious epimutation-selection balance in populations (Day and Bonduriansky, 2011). A key implication of these models—paralleling those involving genetic mutations—is the emergence of ID in consanguineous crosses. Two main features distinguish epimutational from mutational processes. The first is the possibility of paramutation, a process akin to gene conversion in which the epigenetic state of one chromosome can overwrite that of its homolog (Geoghegan and Spencer, 2013). Paramutation tends to reduce epigenetic heterozygosity and may promote the fixation of deleterious epimutants, thereby limiting the expression of ID. The second feature concerns the rates: epimutations often occur at much higher frequencies than genetic mutations and are typically accompanied by a high reversion rate (i.e., resetting to the reference state; Chernomas and Griswold, 2024; Geoghegan and Spencer, 2013). As a result, the contribution of epimutations to ID may be modest in outbred populations, where selection and reversion maintain low equilibrium frequencies. However, in conditions of extreme homozygosity and sustained inbreeding—as in our experimental design—epimutational load could accumulate and regenerate more rapidly than genetic mutations, potentially contributing to the observed fitness decline.

In summary, our study demonstrates that inbreeding depression (ID), typically characterized by reduced offspring fitness with increasing parental relatedness, can appear even in genetically uniform populations. This finding challenges traditional genetic models, as in such contexts individuals cannot differ in genomic heterozygosity. The levels of ID we observed, impacting both juvenile and adult traits, are equivalent to a substantial fraction of the ID typically reported in natural polymorphic populations for the same traits. Although spontaneous mutations continuously arise and create minor genetic differences among individuals, these mutations alone seem insufficient to account for the magnitude of effects we observed across three distinct traits. By definition effects that are not due to genetic differences, should be called epigenetic, irrespective of their material basis. While genetic mutations undoubtedly contribute to ID in populations, it would be premature to assume they account for the entirety of its effects, particularly given the notorious difficulty of fully eliminating ID through purge or fixation. Our findings suggest that epigenetic mechanisms may play a more prominent role in ID than previously recognized––in line with emerging evidence that epimutations and epigenetic inheritance contribute to heritable phenotypic variation in general. Future studies, encompassing a broader range of lines and species, are essential to fully unravel the role and impact of epigenetics to inbreeding depression.

## Methods

### Study organism and experimental conditions

*Physa acuta* is a widespread simultaneously hermaphroditic freshwater snail that reproduces mostly through cross-fertilization. Snails that have no access to mates can resort to self-fertilization (Henry et al., 2005; Jarne et al., 2010). When a potential mate is present, however, individuals nearly always cross-fertilize (Noël et al., 2016b; Pélissié et al., 2012). These snails mate frequently, with unilateral copulations (*i*. *e*., they adopt either the male or the female sex role during a given copulation (Wethington and Dillon, 1996), and can store sperm for long periods. Sexual maturity is reached in around 6 weeks under laboratory conditions at 25 °C.

Throughout the experiments––and also during the establishment of experimental lines––snails were maintained under standard laboratory conditions (25 °C, a 12:12 photoperiod, water renewed twice a week, and *ad*-*libitum* food in the form of boiled ground lettuce). Each individual was kept in a 75 mL plastic box. In addition, albino lines of *P.acuta* are maintained in our laboratory. Albinism (a single-locus recessive trait), can be used to assess paternity success of wild-type focal individuals because wild-type offspring produced by albino mothers must have a wild-type father.

### Field sampling and breeding design

The first step was to collect snails from the field and generate two inbred lines through continuous self-fertilization over many generations. Self-fertilization was enforced by keeping mature individuals isolated until they lay eggs, usually two or three weeks after the age at which they normally start laying eggs when they have mates available (Tsitrone et al., 2003). The two ancestral wild-caught snails were part of a larger sample taken from a wild population at the “Pont Romain” in the Lirou river (43°43’47.0“N, 3°49’50.4”E), located close to the village of Les Matelles, 15 km north of Montpellier (France) on October 23, 2013. The initiation of 44 inbred lines is described in Janicke et al. (2022), who used them for an experiment after the third generation of self-fertilization. A subset of the lines were propagated by continuous self-fertilization until May 2019, a time at which they had all reached at least 25 generations of selfing. The inbreeding coefficient at these stage is therefore expected to be, at least, 1-1/2^25^ = 0.99999997. To evaluate whether inbred individuals were still capable of cross fertilization, we performed a preliminary analysis described in Supporting Information. Once confirmed that individuals from inbred lines were fully capable of performing cross-fertilization, we conducted the main assay.

### Estimating inbreeding depression in outbred clones

We selected two inbred lines named 120 and 110. At this stage, these lines had undergone 28 and 29 generations of self-fertilization, respectively. The residual heterozygosity is expected to be < 4.10^-9^ times that of the wild-caught individuals, ignoring recent mutations. As a first approximation, we therefore consider inbred lines as fully-homozygous clones. The pedigree and the mating protocol described below are depicted in figure 1 and in figures S1 to S6 from the appendix.

### First step: obtaining of F_2_ snails

One individual from each inbred line, 120 and 110, was isolated and forced to self-fertilize to produce the F_1_. F_1_ offspring were isolated at day 28 after egg-laying (still immature) and then raised in isolation until maturity for three more weeks. We assigned a code letter to the F_1_ siblings for identification (120_a_, 120_b_, 120_c_, and so on; and, 110_a_, 110_b_, 110_c_, and so on), constituted inter-line pairs of F_1_ individuals (e.g., 120_a_ x 110_a,_ 120_b_ x 110_b_), and let them mate for one week. To differentiate the 120 snail from its 110 partner in each pair, we added small dots of harmless car-body paint of different colors on their shell (Henry and Jarne, 2007). After mating, F_1_ individuals were re-isolated to lay F_2_ eggs. We successfully collected and raised the resulting F_2_ hatchlings for 10 families *a* to *j* (Fig. 1).

F_2_ snails belonged to two different types, hereafter type S and type T. The S-type F_2_ offspring were laid by a mother from the 120 line, and sired by fathers from the 110 line, while T-type offspring came from the reciprocal combination (mother 110 – father 120). As these two types are present within all 10 families *a* to *j*, we end up with 20 categories (S*_a_*_-*j*_ and T*_a_*_-*j*_), each represented by several offspring. Note that irrespective of the category they belonged to, all F_2_ were expected to be heterozygous and genetically identical to one another (i.e. they form a heterozygous clone; Fig. 1 and S1). We proceeded until we obtained at least 40 F_2_ descendants of type S and 10 of type T for each family. This procedure sometimes required taking several successive clutches or even re-mating the same pair to stimulate egg-laying. These descendants were raised and isolated before maturity, as previously mentioned, and kept until fully mature and ready for mating. Then, we used them to generate different types of F_3_ snails.

### Second step: self-fertilization, sib-, and cousin pair-crosses

We submitted each F_2_ snail to one of three mating treatments: (i) selfing, (ii) sib-cross, and (iii) cousin-cross. Sib and cousin crosses were themselves divided in two subtypes depending on direction of exchange of genetic material (one-and two-ways), giving five treatments in total. We explain this in more detail in the following.

The first treatment included S-type individuals that were forced to self-fertilize (Selfing, Figs. 1 and S2). Thus, ten F_2_ S-type individuals per family (i.e., S*_a_*_,1-10_, S*_b_*_,1-10_, _…_, S*_j_*_,1-10_) were isolated and eggs were collected three days after we detected the first clutch. The second treatment consisted in pair-crossings between two S-type snails from the same family (called “one-way sib cross”). So, for instance, in family *a*, we crossed individuals S_a,11_… S_a,20_, according to the following pairs: S*_a_*_,11/_S*_a_*_,16_, S*_a_*_,12/_S*_a_*_,17_, and so on (and similarly for S*_b_* to S*_j_*) (Figs. 2 and S3). Note that in these crosses the two parents are full sibs and they have the same mother and the same father. The third treatment consisted in pair-crossings within the same family (e.g., family *a*) but between S and T individuals, which we defined as “two-way sib cross”. For instance, we paired snails from S*a* (numbered from 21 to 25) with snails from T*a* (numbered from 1 to 5), which gave us the following combinations: S*_a_*_,21_ / T*_a_*_,1_, S*_a_*_,22_ / T*_a_*_,2_, and S*_a_*_,23_ / T*_a_*_,3_, and so on.

The two parents in these crosses are full-sibs but not obtained in the same way (the father of one is the mother of the other and reciprocally; Figs. 2 and S4). The fourth treatment involved pair-crossings between two F_2_ S-type snails (numbered from 26 to 35) from different families, which we called the one-way cousin cross (the two parents are cousins, belonging to the same type). Because of insufficient individuals in families *h* and *j* we were able to do this for four pairs of families. The resulting combinations were: S*_a_*_,26-35_ / S*_b_*_,26-35_, S*_c_*_,26-35_ / S*_d_*_,26-35_, S*_e_*_,26-35_ / S*_f_*_,26-35_ and S*_g_*_,26-35_ / S*_i_*_,26-35_ (Figs. 2 and S5). Finally, the fifth treatment consisted in pair-crossings between snails of the S-type (36 to 40) and of the T-type (6 to 10) from different families, which we defined as the two-way cousin cross. The combinations were: S*_a_*_,36-40_ / T*_b_*_,6-10_, S*_c_*_,36-40_ / T*_e_*_,6-10_ or S_d,36-40_ / T*_f_*_,6-10_, and so on (Figs. 2 and S6). When necessary, we added different color dots of car-body paint on each individual’s shell to differentiate snails from each family and origin.

After pairing for three days, mated snails were re-isolated to lay eggs for three days. We collected two successive F_3_ clutches for each F_2_ parent, by transferring the parent to a new box after three days (Fig. S9). These F_3_ clutches constitute the generation on which we evaluated fitness traits.

*Third step: assessing and comparing fitness components in F_3_ snails* Eggs were counted in all F_3_ clutches right after collection and live juveniles were counted in the boxes 14 days later. We estimated juvenile survival as the ratio of juveniles over eggs. Next, we randomly selected one of the surviving F_3_ individuals per box and raised it in isolation until age 50 days. At this age we weighed each snail and determined whether it had initiated egg laying or not). Given that each individual was kept in isolation, this egg-laying represents self-fertilization. Then, we paired it with a same-age albino partner for 3 days. Once inseminated, we isolated the F_3_ individual once more, weighed it a second time, and allowed it to lay eggs for 3 days. Finally, we counted these eggs that represent female fecundity of F_3_ under cross-fertilization conditions. We therefore have four traits on the F_3_: juvenile survival, body mass at 50-53 days (the mean of the two measures), self-fertilization ability at 50 days (yes/no), female fecundity after insemination at 54-57 days (number of eggs).

### Heterozygosity

We genotyped a subset of the F_2_ and F_3_ individuals of different categories using 7 microsatellite loci chosen because preliminary data indicated that the inbred lines 110 and 120 were fixed for different alleles. The loci were AF108762, AF108764 (Monsutti and Perrin, 1999), Pac1, Pac2 (Sourrouille et al., 2003), Pasu1-2, Pasu1-9, Pasu1-11 (*cf*. Escobar et al., 2008) grouped into two multiplexes. We expected that the F_2_ would be fully heterozygous (except for a portion of misreadings), whereas in the F_3_ generation, half of the loci would, on average, exhibit heterozygosity. We assessed the number of heterozygous and homozygous loci for each individual. However, it was inevitable that a small fraction of the genotypes remained unamplified, leading to missing data.

Moreover, in microsatellite readings heterozygous genotypes are frequently misconstrued as homozygous, i.e. one allele behaves as partly “dominant” (David et al., 2007). To avoid any bias, we kept the readings blind to the sample’s origin. Also, we refrained from making corrections, even in instances where the reading error was evident (e.g., one homozygous locus and all other loci heterozygous in a F_2_).

### Statistical analyses

All analyses were performed with *R* 4.0.2 packages *lme4* (Bates et al., 2014) and *nlme* (Pinheiro et al., 2017) by fitting generalized linear mixed models (GLMMs) for binomial variables (juvenile survival, ability to self-fertilize, proportion of heterozygous loci) or linear mixed models (LMMs) for Gaussian continuous variables (body weight, female fecundity). Our linear models included as fixed effects: (i) mating treatment (selfing, sib-cross, and cousin-cross), (ii) the cross type (one vs. two ways) when appropriate (i.e., for sib-and cousin-cross), and (iii) the maternal type (S vs. T) within two-way crosses (recall that in the selfing and one-way crosses all mothers are S). We proceeded by first running separate analyses within the sib-mating and cousin-mating treatments in order to test the mating treatment and maternal type effects. As these effects were not significant, we pooled all subcategories within the sib-cross and cousin-cross treatments in order to test the mating treatment effect using the whole dataset. In each model we added an appropriate structure of random effects to account for various levels of replication within each level of fixed factors (e.g., identity of the maternal family: *a*, *b*, *c*; pair of parental families: *a*/*b*, *c*/*d*; identity of the mother: 1 to 35; and/or of the parental pair: 21/1, 22/2 described for each model in the results). Observation number was also added as a random factor in the binomial models on juvenile survival in order to account for overdispersion (Browne et al., 2005; Elston et al., 2001). Tests of fixed effects were made using LRT (Likelihood-ratio tests, which compare the likelihood of models with and without the factor of interest) keeping all random effects present in the model. Random effects were tested using LRT with the corrections indicated by Zuur et al. (2009, chap. 5, pp. 123–125). Post hoc tests were performed when the treatment effect was significant using Holm-Bonferroni correction for multiple-testing to test effect sizes among different treatments.

### Simulations

We made a set of individual-based simulations to evaluate whether the observed changes in fitness between the three type of mating treatments (selfing, sib-cross, cousin-cross) could be compatible with a purely genetic model of inbreeding depression (i.e., deleterious-mutation based). Each individual genome was modelled as a single chromosome on which deleterious mutations were located. The deleterious mutations were of two types: small-effect and large-effect (quasi-lethal), each type with its own rate, degree of recessivity, and effect. We modelled both the process of extraction from a large population and the propagation of two inbred lines over 28 generations with the following events at each generation: mutation, meiosis, recombination, selection, and drift within each line. Then, we simulated the creation of F_1_, F_2_, and the three types of crosses giving the F_3_ as in our experiment. The output of each simulation was a set of three average fitness values for each of the three types of F_3_. For each set of parameter, 1000 independent replicates were made to produce expected distributions of the relative fitnesses of the three categories. The details on the simulations are explained in Support Information in *Simulation Details*.

## Acknowledgments

We thank Tim Janicke for starting the inbred lines. This work was funded by CeMEB (“Projets de Recherche Exploratoires du CeMEB” grant to project “Epigenetics of inbreeding depression (EPID)” by P. David and C. Grunau) and ANR (FRIDA project, grant ANR-23-CE02-0033-01 to P. David). N. Bonel was partially supported by the “Programa de Financiamiento Parcial de Estadías en el Exterior para Investigadores Asistentes”, National Scientific and Technical Research Council CONICET (Res. N° 558 1236/08; 4118/16).

## Authors’ contributions

Conceptualization, CG and PD; methodology, NB and PD; formal analysis, NB and PD; data curation, NB and PD; writing—original draft preparation, NB and PD; writing— review and editing, NB and PD; supervision, PD; project administration, PD; funding acquisition, PD. All authors have read and agreed to the published version of the manuscript.

## References

Agrawal AF, Wang AD. 2008. Increased transmission of mutations by low-condition females: evidence for condition-dependent DNA repair. PLoS Biol 6:e30. doi:10.1371/journal.pbio.0060030

Agrawal AF, Whitlock MC. 2011. Inferences about the distribution of dominance drawn from yeast gene knockout data. Genetics 187:553–566. doi:10.1534/genetics.110.124560

Barrett SCH, Charlesworth D. 1991. Effects of a change in the level of inbreeding on the genetic load. Nature 352:522–524. doi:10.1038/352522a0

Bates D, Mächler M, Bolker B, Walker S. 2014. Fitting Linear Mixed-Effects Models using lme4. arXiv:14065823 [stat].

Biémont C. 2010. Inbreeding effects in the epigenetic era. Nat Rev Genet 11:234–234. doi:10.1038/nrg2664-c1

Bossdorf O, Richards CL, Pigliucci M. 2008. Epigenetics for ecologists. Ecology Letters 11:106–115. doi:10.1111/j.1461-0248.2007.01130.x

Browne WJ, Subramanian SV, Jones K, Goldstein H. 2005. Variance partitioning in multilevel logistics models with over-dispersion. Journal of the Royal Statistical Society 168:599–613. doi:10.1111/j.1467-985X.2004.00365.x

Byers DL, Waller DM. 1999. Do plant populations purge their genetic load? Effects of population size and mating history on inbreeding depression. Annu Rev Ecol Syst 30:479–513. doi:10.1146/annurev.ecolsys.30.1.479

Carr DE, Dudash MR. 1997. The effects of five generations of enforced selfing on potential male and female function in *Mimulus guttatus*. Evolution 51:1797– 1807. doi:10.1111/j.1558-5646.1997.tb05103.x

Cecere G. 2021. Small RNAs in epigenetic inheritance: from mechanisms to trait transmission. FEBS Letters 595:2953–2977. doi:10.1002/1873-3468.14210

Charlesworth B, Borthwick H, Bartolomé C, Pignatelli P. 2004. Estimates of the genomic mutation rate for detrimental alleles in *Drosophila melanogaster*. Genetics 167:815–826. doi:10.1534/genetics.103.025262

Charlesworth B, Charlesworth D. 1999. The genetic basis of inbreeding depression. Genet Res 74:329–340. doi:10.1017/S0016672399004152

Charlesworth B, Charlesworth D. 1998. Some evolutionary consequences of deleterious mutations. Genetica 102/103:3–19. doi:10.1023/A:1017066304739

Charlesworth D, Charlesworth B. 1990. Inbreeding depression with heterozygote advantage and its effect on selection for modifiers changing the outcrossing rate. Evolution 44:870–888. doi:10.1111/j.1558-5646.1990.tb03811.x

Charlesworth D, Willis JH. 2009. The genetics of inbreeding depression. Nat Rev Genet 10:783–796. doi:10.1038/nrg2664

Chelo IM, Afonso B, Carvalho S, Theologidis I, Goy C, Pino-Querido A, Proulx SR, Teotónio H. 2019. Partial Selfing Can Reduce Genetic Loads While Maintaining Diversity During Experimental Evolution. G3 (Bethesda) 9:2811–2821. doi:10.1534/g3.119.400239

Chelo IM, Carvalho S, Roque M, Proulx SR, Teotónio H. 2014. The genetic basis and experimental evolution of inbreeding depression in Caenorhabditis elegans. Heredity 112:248–254. doi:10.1038/hdy.2013.100

Cheptou P-O. 2024. The evolutionary ecology of inbreeding depression in wild plant populations and its impact on plant mating systems. Front Plant Sci 15:1359037. doi:10.3389/fpls.2024.1359037

Cheptou P-O, Donohue K. 2013. Epigenetics as a new avenue for the role of inbreeding depression in evolutionary ecology. Heredity 110:205–206. doi:10.1038/hdy.2012.66

Chernomas G, Griswold CK. 2024. Deleterious mutation/epimutation–selection balance with and without inbreeding: a population (epi)genetics model. GENETICS 227:iyae080. doi:10.1093/genetics/iyae080

Crnokrak P, Barrett SCH. 2002. Perspective: purging the genetic load: a review of the experimental evidence. Evolution 56:2347–2358. doi:10.1111/j.0014-3820.2002.tb00160.x

Dapp M, Reinders J, Bédiée A, Balsera C, Bucher E, Theiler G, Granier C, Paszkowski J. 2015. Heterosis and inbreeding depression of epigenetic Arabidopsis hybrids. Nature Plants 1:15092. doi:10.1038/nplants.2015.92

David P, Pujol B, Viard F, Castella V, Goudet J. 2007. Reliable selfing rate estimates from imperfect population genetic data. Molecular Ecology 16:2474–2487. doi:10.1111/j.1365-294X.2007.03330.x

Day T, Bonduriansky R. 2011. A Unified Approach to the Evolutionary Consequences of Genetic and Nongenetic Inheritance. The American Naturalist 178:E18–E36. doi:10.1086/660911

Elston DA, Moss R, Boulinier T, Arrowsmith C, Lambin X. 2001. Analysis of aggregation, a worked example: numbers of ticks on red grouse chicks. Parasitology 122:563–569. doi:10.1017/S0031182001007740

Escobar JS, Nicot A, David P. 2008. The different sources of variation in inbreeding depression, heterosis and outbreeding depression in a metapopulation of *Physa acuta*. Genetics 180:1593–1608. doi:10.1534/genetics.108.092718

Fallet M, Luquet E, David P, Cosseau C. 2020. Epigenetic inheritance and intergenerational effects in mollusks. Gene 729:144166. doi:10.1016/j.gene.2019.144166

Fry JD, Keightley PD, Heinsohn SL, Nuzhdin SV. 1999. New estimates of the rates and effects of mildly deleterious mutation in *Drosophila melanogaster*. Proc Natl Acad Sci USA 96:574–579. doi:10.1073/pnas.96.2.574

Geoghegan JL, Spencer HG. 2013. The evolutionary potential of paramutation: A population-epigenetic model. Theoretical Population Biology 88:9–19. doi:10.1016/j.tpb.2013.05.003

Glémin S. 2003. How are deleterious mutations purged? Drift versus nonrandom mating. Evolution 57:2678–2687.

Han T, Wang F, Song Q, Ye W, Liu T, Wang L, Chen ZJ. 2021. An epigenetic basis of inbreeding depression in maize. Sci Adv 7:eabg5442. doi:10.1126/sciadv.abg5442

Henry P, Jarne P. 2007. Marking hard-shelled gastropods: tag loss, impact on life-history traits, and perspectives in biology. Invertebrate Biology 126:138–153. doi:10.1111/j.1744-7410.2007.00084.x

Henry P-Y, Bousset L, Sourrouille P, Jarne P. 2005. Partial selfing, ecological disturbance and reproductive assurance in an invasive freshwater snail. Heredity 95:428–436. doi:10.1038/sj.hdy.6800731

Huber CD, Durvasula A, Hancock AM, Lohmueller KE. 2018. Gene expression drives the evolution of dominance. Nat Commun 9:2750. doi:10.1038/s41467-018-05281-7

Janicke T, Chapuis E, Meconcelli S, Bonel N, Delahaie B, David P. 2022. Environmental effects on the genetic architecture of fitness components in a simultaneous hermaphrodite. Journal of Animal Ecology 91:124–137. doi:10.1111/1365-2656.13607

Janicke T, Vellnow N, Sarda V, David P. 2013. Sex-specific inbreeding depression depends on the strength of male-male competition: inbreeding depression in a hermaphrodite. Evolution n/a-n/a. doi:10.1111/evo.12167

Jarne P, Pointier J-P, David P, Koene JM. 2010. Basommatophoran gastropodsThe Evolution of Primary Sexual Characters in Animals. Oxford University Press. pp. 173–196.

Johannes F, Porcher E, Teixeira FK, Saliba-Colombani V, Simon M, Agier N, Bulski A, Albuisson J, Heredia F, Audigier P, Bouchez D, Dillmann C, Guerche P, Hospital F, Colot V. 2009. Assessing the Impact of Transgenerational Epigenetic Variation on Complex Traits. PLoS Genet 5:e1000530. doi:10.1371/journal.pgen.1000530

Kirkpatrick M, Jarne P. 2000. The effects of a bottleneck on inbreeding depression and the genetic load. The American Naturalist 155:154–167. doi:10.1086/303312

Lande R, Schemske DW. 1985. The evolution of self-fertilization and inbreeding depression in plants. I. Genetic models. Evolution 39:24–40. doi:10.1111/j.1558-5646.1985.tb04077.x

Lande R, Schemske DW, Schultz ST. 1994. High inbreeding depression, selective interference among loci, and the threshold selfing rate for purging recessive lethal mutations. Evolution 48:965–978. doi:10.1111/j.1558-5646.1994.tb05286.x

Latter BDH. 1998. Mutant Alleles of Small Effect Are Primarily Responsible for the Loss of Fitness With Slow Inbreeding in Drosophila melanogaster. Genetics 148:1143–1158. doi:10.1093/genetics/148.3.1143

Luquet E, Tariel J. 2016. Offspring reaction norms shaped by parental environment: interaction between within-and trans-generational plasticity of inducible defenses. BMC Evol Biol 16:209. doi:10.1186/s12862-016-0795-9

Luviano N, Lopez M, Gawehns F, Chaparro C, Arimondo PB, Ivanovic S, David P, Verhoeven K, Cosseau C, Grunau C. 2021. The methylome of Biomphalaria glabrata and other mollusks: enduring modification of epigenetic landscape and phenotypic traits by a new DNA methylation inhibitor. Epigenetics & Chromatin 14:48. doi:10.1186/s13072-021-00422-7

McCarthy TM, Sih A. 2008. Relatedness of mates influences mating behaviour and reproductive success of the hermaphroditic freshwater snail Physa gyrina. Evolutionary Ecology Research 10:77–94.

Monsutti A, Perrin N. 1999. Dinucleotide microsatellite loci reveal a high selfing rate in the freshwater snail *Physa acuta*. Molecular Ecology 8:1076–1078. doi:10.1046/j.1365-294X.1999.00655_2.x

Noël E, Chemtob Y, Janicke T, Sarda V, Pélissié B, Jarne P, David P. 2016a. Reduced mate availability leads to evolution of self-fertilization and purging of inbreeding depression in a hermaphrodite. Evolution 70:625–640. doi:10.1111/evo.12886

Noël E, Chemtob Y, Janicke T, Sarda V, Pélissié B, Jarne P, David P. 2016b. Reduced mate availability leads to evolution of self-fertilization and purging of inbreeding depression in a hermaphrodite: selfing evolution in a hermaphroditic snail. Evolution 70:625–640. doi:10.1111/evo.12886

Pélissié B, Jarne P, David P. 2012. Sexual selection without sexual dimorphism: bateman gradients in a simultaneous hermaphrodite: sexual selection in a hermaphrodite. Evolution 66:66–81. doi:10.1111/j.1558-5646.2011.01442.x

Pinheiro J, Bates D, DebRoy S, Sarkar D, R Core Team. 2017. nlme: linear and nonlinear mixed effects models. R package version 3.1–131.

Preite V, Oplaat C, Biere A, Kirschner J, Van Der Putten WH, Verhoeven KJF. 2018. Increased transgenerational epigenetic variation, but not predictable epigenetic variants, after environmental exposure in two apomictic dandelion lineages. Ecology and Evolution 8:3047–3059. doi:10.1002/ece3.3871

Richards EJ. 2006. Inherited epigenetic variation — revisiting soft inheritance. Nat Rev Genet 7:395–401. doi:10.1038/nrg1834

Rumball W, Franklin IR, Frankham R, Sheldon BL. 1994. Decline in heterozygosity under full-sib and double first-cousin inbreeding in Drosophila melanogaster. Genetics 136:1039–1049. doi:10.1093/genetics/136.3.1039

Sharp NP, Agrawal AF. 2012. Evidence for elevated mutation rates in low-quality genotypes. Proc Natl Acad Sci USA 109:6142–6146. doi:10.1073/pnas.1118918109

Simmons MJ, Crow JF. 1977. Mutations affecting fitness in drosophila populations. Annu Rev Genet 11:49–78. doi:10.1146/annurev.ge.11.120177.000405

Sourrouille P, Debain C, Jarne P. 2003. Microsatellite variation in the freshwater snail *Physa acuta*. Molecular Ecology Notes 3:21–23. doi:10.1046/j.1471-8286.2003.00338.x

Tariel J, Plénet S, Luquet É. 2020. Transgenerational plasticity of inducible defences: Combined effects of grand-parental, parental and current environments. Ecology and Evolution 10:2367–2376. doi:10.1002/ece3.6046

Tsitrone A, Duperron S, David P. 2003. Delayed Selfing as an Optimal Mating Strategy in Preferentially Outcrossing Species: Theoretical Analysis of the Optimal Age at First Reproduction in Relation to Mate Availability. The American Naturalist 162:318–331. doi:10.1086/375542

Van Der Graaf A, Wardenaar R, Neumann DA, Taudt A, Shaw RG, Jansen RC, Schmitz RJ, Colomé-Tatché M, Johannes F. 2015. Rate, spectrum, and evolutionary dynamics of spontaneous epimutations. Proc Natl Acad Sci USA 112:6676–6681. doi:10.1073/pnas.1424254112

Vergeer P, Wagemaker N (C. AM), Ouborg NJ. 2012. Evidence for an epigenetic role in inbreeding depression. Biol Lett 8:798–801. doi:10.1098/rsbl.2012.0494

Waller DM. 2021. Addressing Darwin’s dilemma: Can pseudo-overdominance explain persistent inbreeding depression and load? Evolution 75:779–793. doi:10.1111/evo.14189

Wang J, Hill WG. 1999. Effect of selection against deleterious mutations on the decline in heterozygosity at neutral loci in closely inbreeding populations. Genetics 153:1475–1489. doi:10.1093/genetics/153.3.1475

Wethington AR, Dillon, Jr RT. 1996. Gender choice and gender conflict in a non-reciprocally mating simultaneous hermaphrodite, the freshwater snail,Physa. Animal Behaviour 51:1107–1118. doi:10.1006/anbe.1996.0112

Williams JL, Hall SJG, Del Corvo M, Ballingall KT, Colli L, Ajmone Marsan P, Biscarini F. 2016. Inbreeding and purging at the genomic Level: the Chillingham cattle reveal extensive, non-random SNP heterozygosity. Animal Genetics 47:19–27. doi:10.1111/age.12376

Zhao L, Charlesworth B. 2016. Resolving the Conflict Between Associative Overdominance and Background Selection. Genetics 203:1315–1334. doi:10.1534/genetics.116.188912

Zuur AF, Ieno EN, Walker N, Saveliev AA, Smith GM. 2009. Mixed effects models and extensions in ecology with R, Statistics for Biology and Health. New York, NY: Springer New York. doi:10.1007/978-0-387-87458-6

