## Supporting Information for "Not just mutations: Inbreeding depression persists without genetic variation"

#### Table of Contents


***Supplementary Methods and Results***

Preliminary assay: Outcrossing ability of inbred lines 2

Simulations 2

References 6

***Supplementary Tables***

*Table S1*. Results of linear models for fitness traits: all treatments pooled 7

*Table S2*. Results of linear models for fitness traits: sib-cross 9

*Table S3*. Results of linear models for fitness traits: cousin-cross 10

***Supplementary Figures***

*Figure S1*. Pedigree of the $F_2$ snails: experimental strains 11

*Figure S2*. Pedigree of $F_3$ from the *Selfing* treatment 12

*Figure S3*. Pedigree of $F_3$ from the *Sib-cross* treatment: one way 13

*Figure S4*. Pedigree of $F_3$ from the *Sib-cross* treatment: two ways 14

*Figure S5*. Pedigree of $F_3$ from the *Cousin-cross* treatment: one way 15

*Figure S6*. Pedigree of $F_3$ from the *Cousin-cross* treatment: two ways 16

*Figure S7*. Results of simulations with different parameter sets 17

*Figure S8*. Results of simulations for increased mutation rates 18

### Supplementary Methods and Results

#### Preliminary assay to evaluate the outcrossing ability of inbred lines

We conducted a preliminary assay to confirm whether inbred individuals were still capable of outcrossing considering that they were forced to self-fertilization, and had not practiced sexual behavior for more than 25 generations. This assay was split in two steps. The first one consisted in coupling three to four virgin individuals (focals) from each of nine inbred lines (named 45, 46, 85, 99, 101, 108, 110, 111, 120) with a virgin mature albino partner (outcrossing) and let them in pairs for three days. After the 3-day pairing period, focals and albinos were isolated for five (5) days to lay eggs. We let these egg hatch and raised the progeny. Juveniles emerging from the eggs laid by the albino mate were pigmented, an indicator of the ability of inbred individuals to cross-fertilize eggs of their partners. The progeny obtained from the eggs laid by the inbred individual was all pigmented as the pigmented allele is dominant. To check that the progeny were heterozygous (and therefore that inbred individuals let their own eggs be cross-fertilized by a partner) we grew some of them to adulthood and test-crossed them by again mating them with albino. The overall proportion of pigmented offspring from albino partners was 0.42±0.05 SE (n = 22, excluding zeroes). The presence of albinos in the progeny of inbred lines indicated that the tested individuals were indeed heterozygous, which confirmed that inbred individuals were fully capable of performing cross-fertilization even after more than 25 generations of inbreeding.

#### Simulation details

The simulations were individual-based, each individual was represented by a diploid genome, consisting of a single chromosome pair. For simplicity we preferred to manipulate the degree of linkage through the length of a single chromosome rather than the number of chromosomes. Each chromosome was itself represented by two lists of numbers, representing the positions of two types of deleterious mutations (semi lethal-effect [*d1*] and small-effect [*d2*]) along the chromosome. The lists were of variable length depending on the number of mutations in each genome, which varied across individuals. The parameters input before each simulation were: the effects *s1, s2* and dominance *h1, h2* of the two types of mutations, the fitness effects (1-*s* and 1-*hs* for homozygous and heterozygous carriers of the mutation, respectively), the amount of inbreeding depression (relative decrease in average fitness between selfed and outcrossed offspring) contributed by the first and second type of deleterious mutations (*d1* and *d2*) in the initial founder population, and the total length (L) of the genome (in morgans).

Each simulation consisted in three steps (i) computing two genomic mutation rates *U1* and *U2* (one for each type of mutation) compatible with the input parameters (ii) simulating the extraction and propagation of inbred (selfing) lines over 28 generations of mutation-recombination-selection-drift (iii) simulating the $F_1$, $F_2$ and $F_3$ obtained by crossing two independent lines and computing their average fitness.

<u>Step 1:</u> *Obtaining of mutation rates compatible with input parameters*

We considered that the original population was near-infinite so that the frequency of deleterious mutations at each locus was the mutation-selection equilibrium under random mating. The usual approximations are $u/hs$ for non-fully recessive mutations (*h*>0) and $\sqrt{u/s}$ for fully recessive ones (*h*=0), with *u* the per-locus deleterious mutation rate (Crow and Kimura, 1970). Because we explored very low values for *h* we did not make these approximations and instead used the exact solution of the (quadratic) equilibrium equation which is:

$$q = \frac{-hs(1+2u)+\sqrt{(hs(1+2u))^2+4us(1-2h)(1+u)}}{2s(1+2hs(+u)}$$

The two expressions above are limit cases of this solution. The total inbreeding depression *d* generated by *n* such loci adding multiplicatively is $1 - Exp(-ns\left(\frac{1}{2}-h\right)q)$ yielding $q = -\frac{Log(1-d)}{\left(\frac{1}{2}-h\right)sn}$. By equating the two expressions of *q* and solving for *u* one can find the per-locus mutation rate (and, multiplying by *n*, the per-haploid genome mutation rate *U*) necessary to generate the observed inbreeding depression *d* given selective coefficients (*s*) and dominance (*h*). We used *n*=10,000 as the number of mutable genes in the genome but, in practice, this number had very little effect on the resulting *U* provided it was not very small (*i.e.* <100). We did these computations separately for each category of mutation, yielding *U1* and *U2*, the corresponding genomic mutation rates,as a function of *d1* and *d2*, the ID generated by large- and small-effect mutations respectively. In the simulations the sum of *d1* and *d2* was constrained to equal the average ID for juvenile survival measured in the laboratory on the progeny of wild-caught snails from natural populations in the Montpellier region, where our ancestral snails were collected (Escobar et al., 2008; Janicke et al., 2013; see "parameters explored" section below).

Step 2: *Simulation of inbred lines*

The expected number of mutations per haploid genome in the panmictic population of origin is *nq*. We initiated each line by drawing two random numbers in a Poisson distribution of mean *nq* representing mutations carried by the paternal and maternal chromosomes of the founding individual. We assumed that all mutations were initially heterozygous and drew their positions at random in a Uniform distribution (minimum 0, maximum L, the length of the chromosome).

At each generation within an inbred line, all offspring are produced by self-fertilization of a single individual. Ideally only one of the offspring is grown to adulthood and serves to produce the new generation. In practice, however, self-fertilized broods are made of several juveniles, which represent survivors of a survival selection process (e. g. after elimination of homozygous lethals), and, among these juveniles, we grow not only one but a few to adulthood. Having several adults provides backups in case one dies without reproducing or fails to reproduce. This means that, in practice, the size of the adult population before selection is not 1 but, in our case, around N=3 within each line. For simplification, we assumed that all semi lethal-effect mutations (type 1) acted first at an early stage while small-effect mutations (type 2) acted at a later stage on the survivors of the previous stage. That is, the fitness components *w1* (first stage) and *w2* (second stage) were acting in sequence in the simulated passage of generations. The successive steps to create a new generation, starting from N adults, were the following:

a) Eliminate all fixed mutations (mutations homozygous in all *N* adults) as they will have no effects on relative fitness and selection within the line.
b) Select one parent among N with probability proportional to *w2*.
c) Create two gametes from the selected parent. Each gamete is produced by first drawing a recombination number *r* (Poisson-distributed with mean L), then randomly placing *r* recombination breakpoints in the map (uniform distribution between 0 and L), then reassembling bits of the paternal and maternal chromosomes according to recombination breakpoints, then adding Poisson-distributed numbers of neo-mutations of type 1 and 2 (means $U_1$ and $U_2$), uniformly distributed along the genome.
d) Compute the fitness *w1* and *w2* for the zygote made by the union of the two gametes (multiplicative fitness over loci); keep the zygote with probability *w1*.
e) Repeat steps *b*, *c* and *d* until N zygotes are conserved. They become the adults of the next generation.

This process was repeated over 28 generations to mimic the creation of our inbred lines.

Step 3: *Simulation of the experiment*

In this step we randomly drew one individual from generation 28 in one line, and propagated it by self-fertilization with recombination, mutation and selection as above to produce 10 $F_1$ offspring. We then simulated the mating of these 10 offspring to 10 corresponding offspring of another (independently constituted) line resulting in 10 sets of two $F_2$. For each mating, the procedure was the same as in the passage of generations (*cf.* above) except that zygotes were constituted by drawing gametes from different parents. Finally, we computed the average fitness of $F_3$ for different types of matings: self-fertilization of $F_2$, sib-mating of $F_2$ (mating the two $F_2$ produced by the same pair of $F_1$), cousin-mating of $F_2$ (mating between two $F_2$ from different pairs). For each type of mating we computed the fitnesses *w1* and *w2* and their product *w* in 100 independent zygotes made with recombination and mutation, but not selection from the chosen parents (as we want the fitnesses before selection), and averaged over the 100 different zygotes. For each simulated pair of lines, we computed the ratio of average fitnesses between offspring of cousin-cross and selfing, and offspring of sib-cross and selfing, as in our experiment. Then, we simulated 1000 independent line pairs for each parameter set and presented the means and SD of fitness ratios over the 1000 replicates. The simulations were made using a Mathematica 5.0 program available on request.

Parameters explored:

Among the traits studied in our experiment, the one for which inbreeding depression is the largest in natural populations is juvenile survival, with an average around *d*=0.5 (Escobar et al., 2008; Janicke et al., 2013). We used this value as a starting point; our reference situation is *d1*=0.3 and *d2*=0.2 (depression contributed by semi lethal and weak-effect mutations, respectively); *s1*=0.9 and *h1*=0.02 for semi lethal-effect; *s2*=0.05 and *h2*=0.2 for small-effect mutations, and a genome length of *L* =10 morgans (probably underestimated as *P. acuta* has 18 pairs of chromosomes but the results are insensitive to further increases in *L*). In the reference situation, the per-haploid genome mutation rates are $U_1$ = 0.015 and $U_2$ = 0.15 per genome, resulting in an average load of 0.82 semi lethal and 14.9 small-effect deleterious mutations per gamete in the initial population. Recall that these are not all the mutations but only the ones affecting one particular fitness component, juvenile survival; results are therefore to be compared to those obtained on juvenile survival in our experiment. Traits expressed later contribute less to ID in snails from natural populations, and so would require lowering ID (to values <0.5), and therefore mutation rates and/or mutational effects (s). For this reason we consider that parameter sets based on juvenile survival generate maximal expectations for ID on any particular trait. We explored various deviations from the reference situation (but still keeping it compatible with a total ID=0.5 in natural populations) by changing the relative weights of semi lethal- versus small-effect mutations ($d_1$, $d_2$ = 0.49, 0.01; 0.4, 0.1; 0.3, 0.2; 0.2, 0.3; 0.1, 0.4; 0.01, 0.49); the recessivity and strength of semi lethal-effect ($h_1$=0.01, 0.02, 0.05; $s_1$= 0.8, 0.9, 0.99) and of weak-effect mutations ($h_2$ = 0.1, 0.2, 0.4; $s_2$ = 0.025, 0.05, 0.1), and genome length ($L$ = 10, 5, 1), see Fig. S7. Note that very tight linkage ($L$ = 1) is unrealistic but this is the scenario that may maximize the constitution of non-recombining blocks of mutations, resulting in pseudo-overdominance and persistence of heterozygosity within the lines (Waller, 2021). Finally, we explored situations with the mutation rates increased up to 15 times the reference values (and therefore not compatible with available estimates of ID in natural populations), to evaluate how much mutation rates should be increased to generate results matching our observations (see Fig S8).

#### References


Crow JF, Kimura M. 1970. An introduction to population genetics theory. New Jersey: Blackburn Press.

Escobar JS, Nicot A, David P. 2008. The different sources of variation in inbreeding depression, heterosis and outbreeding depression in a metapopulation of *Physa acuta*. Genetics 180:1593–1608. doi:10.1534/genetics.108.092718

Janicke T, Vellnow N, Sarda V, David P. 2013. Sex-specific inbreeding depression depends on the strength of male-male competition: inbreeding depression in a hermaphrodite. Evolution doi:10.1111/evo.12167

Waller DM. 2021. Addressing Darwin's dilemma: Can pseudo-overdominance explain persistent inbreeding depression and load? Evolution 75:779–793. doi:10.1111/evo.14189

#### Supplementary Tables

**Table S1.** Detailed results of the linear models on fitness traits measured on $F_3$ offspring resulting from different mating treatments (selfing, sib- and cousin-cross) of the hermaphrodite snail *Physa acuta*. Significant values of each effect are indicated in bold. In the two pair-cross treatments there are two possible maternal types (S versus T) but the difference between them was never significant (see Tables S2, S3). For simplicity, we therefore ignored Maternal type in this analysis. Similarly the pair-cross treatments include two categories (one-way *vs*. two-way). The difference between them was significant only for juvenile survival in the sib-cross treatment and fecundity rate in the cousin-cross. We therefore provide tests of mating treatment effects either pooling one-way and two-way categories within the sib-cross and cousin-cross treatments (top) or reducing the dataset to the one-way category (bottom). The results are similar. The random effects (nested within treatments) include (i) the parental family: the letters or pair of letters characterizing the origin of the $F_2$ parents used (*selfing*: a, b, c, where there is only one parent; *sib-cross*: aa, bb, cc, where there are two parents with the same letter; *cousin-cross*: ab, cd, ef, where there are two parents with different letters). (ii) The parental couple: nested within the parental family, labelled with different numbers (e.g. 1, 2, 3, etc.). For the *selfing* treatment we considered that the pair was made of the individual with itself. (iii) The maternal individual: nested within the parental couple. This component of variance has two possibilities in each couple and is estimated only based on pair-crosses, as by definition, the parental couple and the mother are confounded in the *selfing* treatment. (iv) An “observation” factor takes a different value for each observation and allows for considering overdispersion, if present, in the binomial models. Recall that there are several observations per mother (e.g. two successive clutches in which we counted juvenile survival, several $F_3$ individuals kept to adulthood.)

| | Variable | Fixed effects: Mating treatment | Random effects: parental family | parental couple | mother | individual | Model | Number of observations |
|---|---|---|---|---|---|---|---|---|
| **All mating treatments** | Juvenile Survival | $X^2_2 = 15.40$ *P* **<0.001** | variance = 0.10 *P* **<0.001** | variance = 0 | variance = 0.01 *P* = 0.299 | variance = 0.84 *P* **<0.001** | Binomial | 439 |
| | Probability of selfing | $X^2_2 = 19.69$ *P* **<0.001** | variance = 0.26 *P* **= 0.033** | variance = 0 | variance = 0.11 *P* = 0.228 | variance = 0 | Binomial | 368 |
| | Body weight | $X^2_2 = 10.06$ *P* **= 0.007** | variance = 68.15 *P* **<0.001** | variance = 0 | variance = 0 | | Gaussian | 368 |
| | Fecundity rate | $X^2_2 = 1.53$ *P* = 0.466 | variance = 0 | variance = 0 | variance = 0.20 *P* = 0.375 | | Gaussian | 353 |
| All mating treatments (but only 1 way) | Juvenile Survival | $X^2_2 = 13.93$ *P* **<0.001** | variance = 0.07 *P* **= 0.010** | variance = 0 | variance = 0 | variance = 0.82 *P* **<0.001** | Binomial | 268 |
| | Probability of selfing | $X^2_2 = 12.57$ *P* **= 0.002** | variance = 0.52 *P* **= 0.007** | variance = 0 | variance = 0.23 *P* = 0.244 | variance = 0 | Binomial | 226 |
| | Body weight | $X^2_2 = 7.38$ *P* **= 0.025** | variance = 95.02 *P* **<0.001** | variance = 0 | variance = 0 | | Gaussian | 226 |
| | Fecundity rate | $X^2_2 = 1.51$ *P* = 0.470 | variance = 0 | variance = 0 | variance = 0.17 | | Gaussian | 219 |

**Table S2.** Detailed results of the linear models on fitness traits measured on $F_3$ offspring resulting from different sibling pair crossing testing the effect of direction (1 way [1w] and 2 ways [2w]), and the effect of parental origin (S and T) within the 2w category (where each pair has one T and one S parent) . Significant values of each effect are indicated in bold. The random effects are the same as previously plus some interaction with the fixed effects tested, when appropriate.

| | Variable | Fixed effects: Direction (1 way vs. 2 way) | Fixed effects: Origin (S vs. T) | Random effects: parental family | Random effects: parental couple | Random effects: mother | Random effects: mother family | Random effects: individual | Random effects: Origin \| parental family | Random effects: Origin \| parental couple | Model | Number of observations |
|---|---|---|---|---|---|---|---|---|---|---|---|---|
| Cousins (1w vs. 2w) | Juvenile Survival | $X^2_1$ = 1.92<br>*P* = 0.589 | | variance = 0.10<br>*P* = 0.073 | variance = 0.02<br>*P* = 0.5 | variance = 0.02<br>*P* = 0.5 | variance = 0.07<br>*P* = 0.180 | variance = 0.71<br>*P* **<0.001** | | | Binomial | 153 |
| | Probability of selfing | $X^2_1$ = 2.74<br>*P* = 0.098 | | variance = 0.22<br>*P* = 0.214 | variance = 0.16<br>*P* = 0.5 | variance = 0.20<br>*P* = 0.5 | | variance = 0 | | | Binomial | 134 |
| | Body weigh | $X^2_1$ = 0.003<br>*P* = 0.954 | | variance = 0.46<br>*P* **= 0.003** | variance = 0.60<br>*P* = 0.5 | variance = 0 | | | | | Gaussian | 134 |
| | Fecundity rate | $X^2_1$ = 6.27<br>*P* = 0.014 | | variance = 0 | variance = 0 | variance = 0 | | | | | Gaussian | 128 |
| Cousins (S vs. T) | Juvenile Survival | | $X^2_1$ = 0.02<br>*P* = 0.875 | | | | | variance = 0.88<br>*P* **<0.001** | var. par. family = 0.02<br>var. Origin * family = 0.55<br>*P* = 0.088 | var. par. couple = 0.21<br>var. Origin * couple= 0.21<br>*P* = 0.265 | Binomial | 73 |
| | Probability of selfing | | $X^2_1$ = 0.72<br>*P* = 0.131 | | | | | variance = 0 | var. par. family = 0<br>var. Origin * family = 1.27<br>*P* = 0.300 | var. par. couple = 0<br>var. Origin * couple= 0.1 | Binomial | 63 |
| | Body weight | | $X^2_1$ = 3.53<br>*P* = 0.060 | | | | | | var. par. family = 30.41<br>var. Origin * family = 72.61<br>*P* = 0.370 | var. par. couple = 0<br>var. Origin * couple = 6.18 | Gaussian | 63 |
| | Fecundity rate | | $X^2_1$ = 0.02<br>*P* = 0.883 | | | | | | var. par. family = 7.90<br>var. Origin * family = 9.17<br>*P* = 0.087 | var. par. couple = 0<br>var. Origin * couple= 0<br>*P* = 0.49 | Gaussian | 58 |

**Table S3.** Detailed results of the linear models on fitness traits measured on $F_3$ offspring resulting from different cousin pair crossing testing the effect of direction (1 way [1w] and 2 ways [2w]) and the effect of origin (S and T) within 2w crosses, in *Physa acuta*. Significant values of each effect are indicated in bold.

| | Variable | Fixed effects: Direction (1 way vs. 2 way) | Fixed effects: Origin (S vs. T) | Random effects: parental family | parental couple | mother | mother family | observation | Origin \| parental family | Origin \| parental couple | Model | Number of observations |
|---|---|---|---|---|---|---|---|---|---|---|---|---|
| Cousins (1w vs. 2w) | Juvenile Survival | $X^2_1$ = 1.92 | | variance = 0.10 | variance = 0.02 | variance = 0.02 | variance = 0.07 | variance = 0.71 | | | Binomial | 153 |
| | | *P* = 0.589 | | *P* = 0.073 | *P* = 0.5 | *P* = 0.5 | *P* = 0.180 | *P* **<0.001** | | | | |
| | Probability of selfing | $X^2_1$ = 2.74 | | variance = 0.22 | variance = 0.16 | variance = 0.20 | | variance = 0 | | | Binomial | 134 |
| | | *P* = 0.098 | | *P* = 0.214 | *P* = 0.5 | *P* = 0.5 | | | | | | |
| | Body weigl | $X^2_1$ = 0.003 | | variance = 0.46 | variance = 0.60 | variance = 0 | | | | | Gaussian | 134 |
| | | *P* = 0.954 | | *P* **= 0.003** | | | | | | | | |
| | Fecundity rate | $X^2_1$ = 6.27 | | variance = 0 | variance = 0 | variance = 0 | | | | | Gaussian | 128 |
| | | *P* **= 0.014** | | | | | | | | | | |
| Cousins (S vs. T) | Juvenile Survival | | $X^2_1$ = 0.02 | | | | | variance = 0.88 | var. par. family = 0.02 | var. par. couple = 0.21 | Binomial | 73 |
| | | | *P* = 0.875 | | | | | *P* **<0.001** | var. Origin * family = 0.55 | var. Origin * couple = 0.21 | | |
| | | | | | | | | | *P* = 0.088 | *P* = 0.265 | | |
| | Probability of selfing | | $X^2_1$ = 0.72 | | | | | variance = 0 | var. par. family = 0 | var. par. couple = 0 | Binomial | 63 |
| | | | *P* = 0.131 | | | | | | var. Origin * family = 1.27 | var. Origin * couple = 0.1 | | |
| | | | | | | | | | *P* = 0.300 | *P* = 0.5 | | |
| | Body weight | | $X^2_1$ = 3.53 | | | | | | var. par. family = 30.41 | var. par. couple = 0 | Gaussian | 63 |
| | | | *P* = 0.060 | | | | | | var. Origin * family = 72.61 | var. Origin * couple = 6.18 | | |
| | | | | | | | | | *P* = 0.370 | *P* = 0.5 | | |
| | Fecundity rate | | $X^2_1$ = 0.02 | | | | | | var. par. family = 7.90 | var. par. couple = 0 | Gaussian | 58 |
| | | | *P* = 0.883 | | | | | | var. Origin * family = 9.17 | var. Origin * couple = 0 | | |
| | | | | | | | | | *P* = 0.087 | *P* = 0.49 | | |

#### Supplementary Figures

**Figure S1.** Pedigree of the $F_2$ snails used in the mating treatment approach of this study. Parents were constituted by inbred lines (120 and 110) that had undergone more than 25 generations of self-fertilization prior to the experiment. These lines are considered purely homozygous, neglecting recent mutations, as the fraction of the initial heterozygosity that remains in their genome ($0.5^{28\ \text{generations}}$) is lower than $4e^{-9}$. Green and blue bars indicate the line of origin of the chromosomes. $F_1$ individuals (inbred sublines) resulted from self-fertilization of parents, i.e. they have the same genetic architecture. The $F_1$ pair crossing yielded a set of individuals that constituted the experimental strains ($F_2$ snails), which are all heterozygous and clonal to one another but with different type of origins, S and T. $F_2$ snails from the S type had their mothers from the 120 line and their fathers from the 110 line, and vice versa for those from type T. Black dots depict zygotes and red and black lines or arrows represent male and female meiotic cycles, respectively.

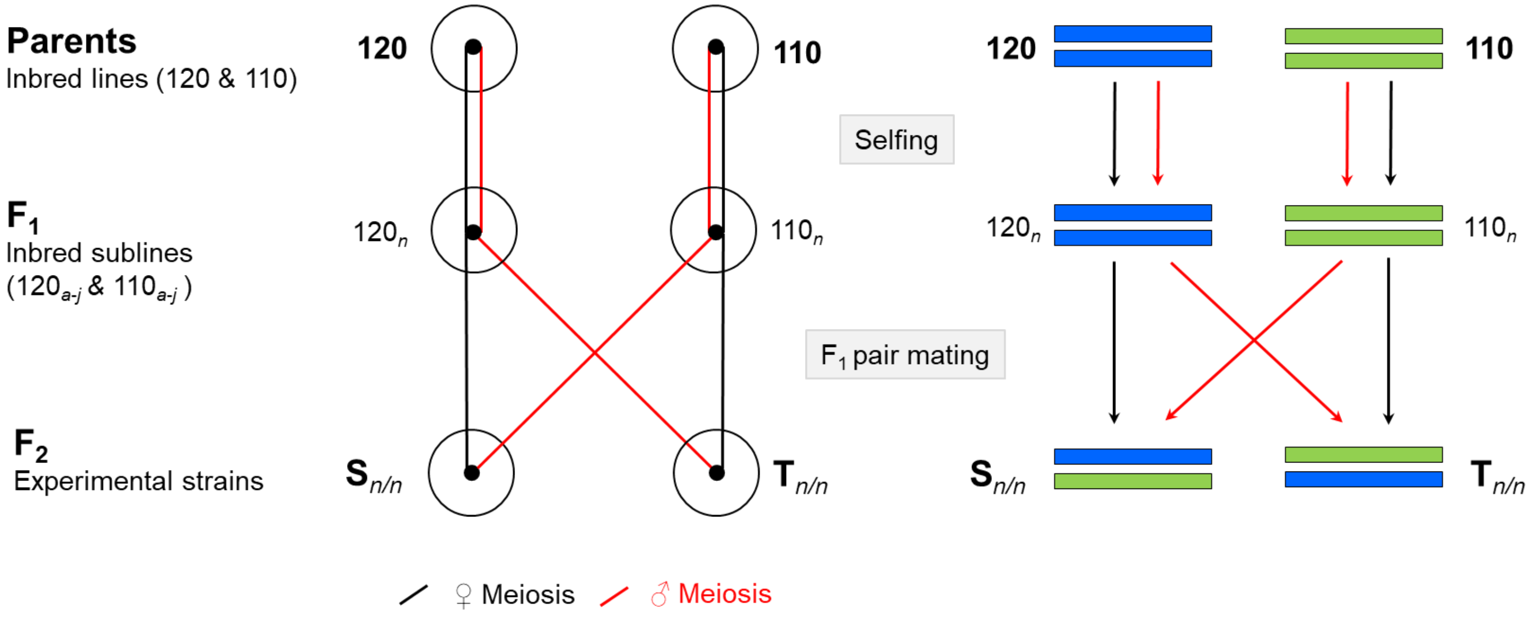

**Figure S2.** Pedigree of $F_3$ snails resulting from self-fertilization (Selfing) of a $F_2$ individuals from the S origin. $F_3$ snails have fragments of chromosomes that are homozygous by descent (or identical by descent, *ibd*). Each homozygous region (e.g. blue or green region surrounded by a dashed rectangle) is composed of two fragments copied from the same original chromosome. The homozygous regions represent 50% of the genome. Within these regions the distance between the maternal and paternal fragments is two (i.e. number of meiotic cycles from their common ancestral chromosome). Each of the two gametes has undergone one meiotic cycle, one during male gametogenesis and the other during female gametogenesis (one possible pathway is highlighted in yellow).

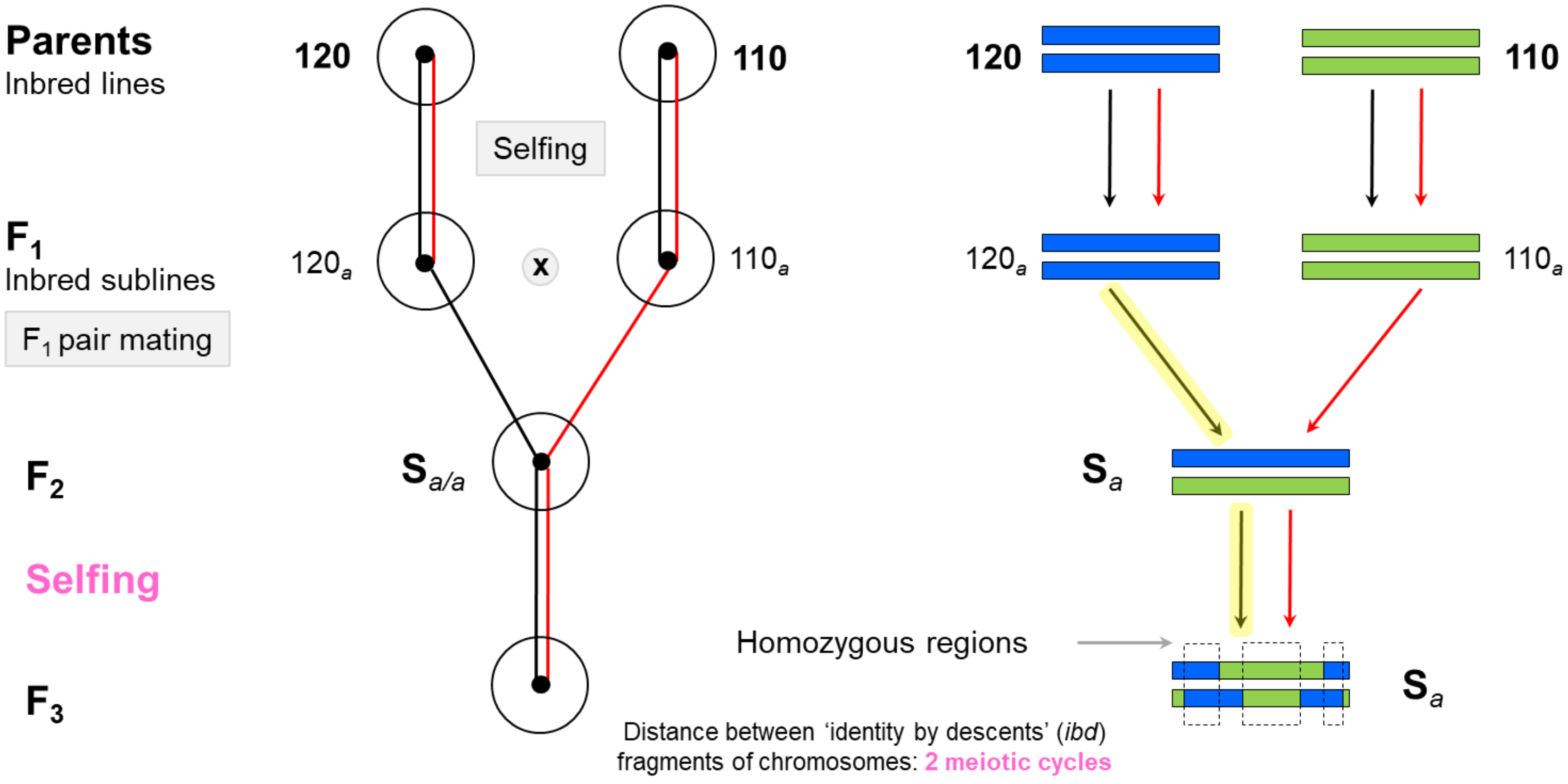

**Figure S3.** Pedigree of $F_3$ snails resulting from a one-way sib-cross between $F_2$ individuals within the S origin and within the same family (e.g., *a*), which are genetically identical to each other. The distance between the *ibd* fragments to coalescence to the ancestral chromosome ($F_1$) is at least four meiotic cycles (path highlighted in yellow). This is, in *ibd* chromosomal regions for the "120" alleles (blue) at least three female meiotic cycles (black segments) and one male meiotic cycle (red segments) separate the maternal and paternal copy (one possible pathway is highlighted in yellow). In *ibd* chromosomal regions for the "110" alleles (green), it is at least three male meiotic cycles and one female meiotic cycle.

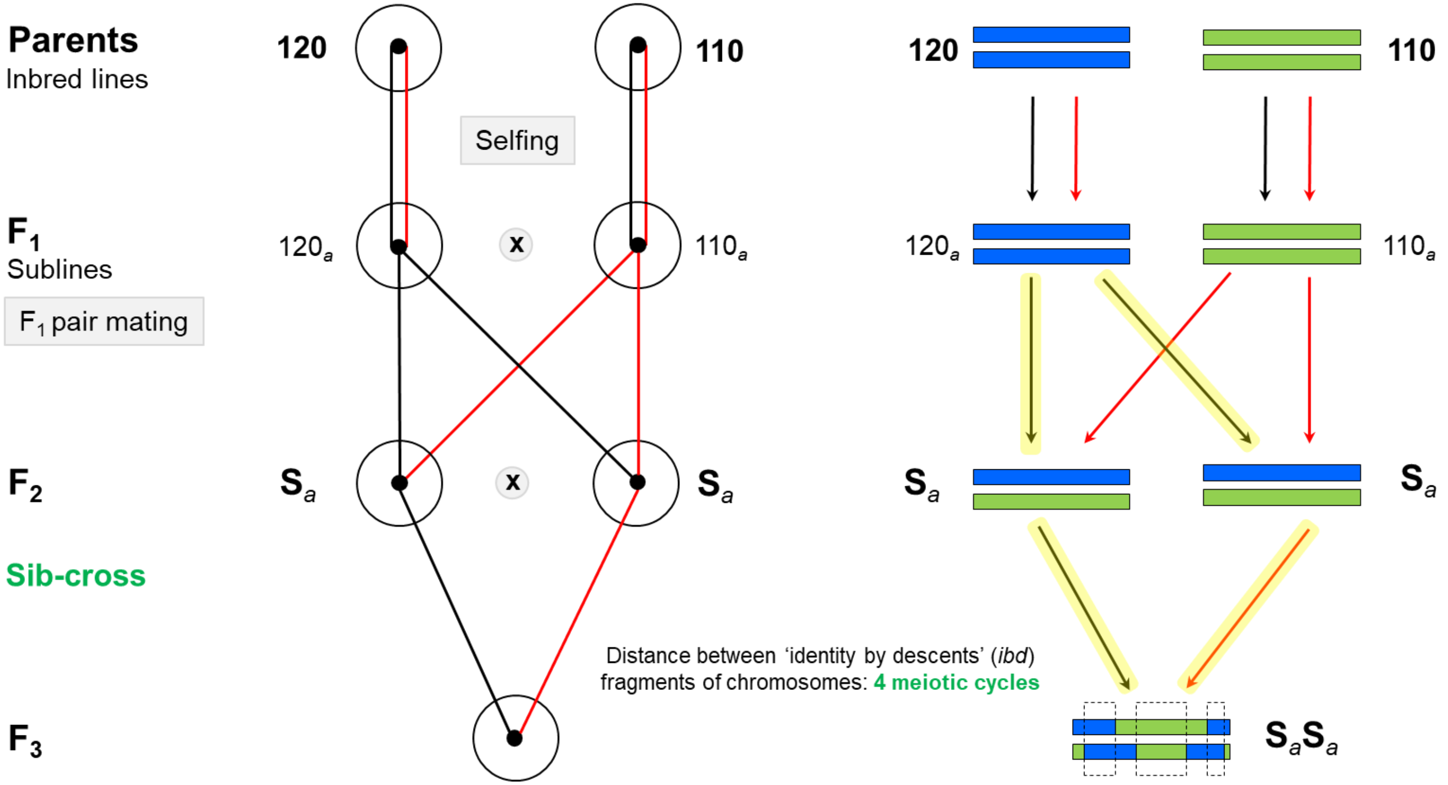

**Figure S4.** Pedigree of $F_3$ snails resulting from a two-way sib-cross between $F_2$ individuals from different origins (S and T) but from the same family (e.g., *a*). The distance between the *ibd* fragments to coalescence to the ancestral chromosome is at least four meiotic cycles, two male and two female meiotic cycles, irrespective of whether the *ibd* region is of 120 or 110 origin (i.e. blue or green, respectively; one possible pathway is highlighted in yellow).

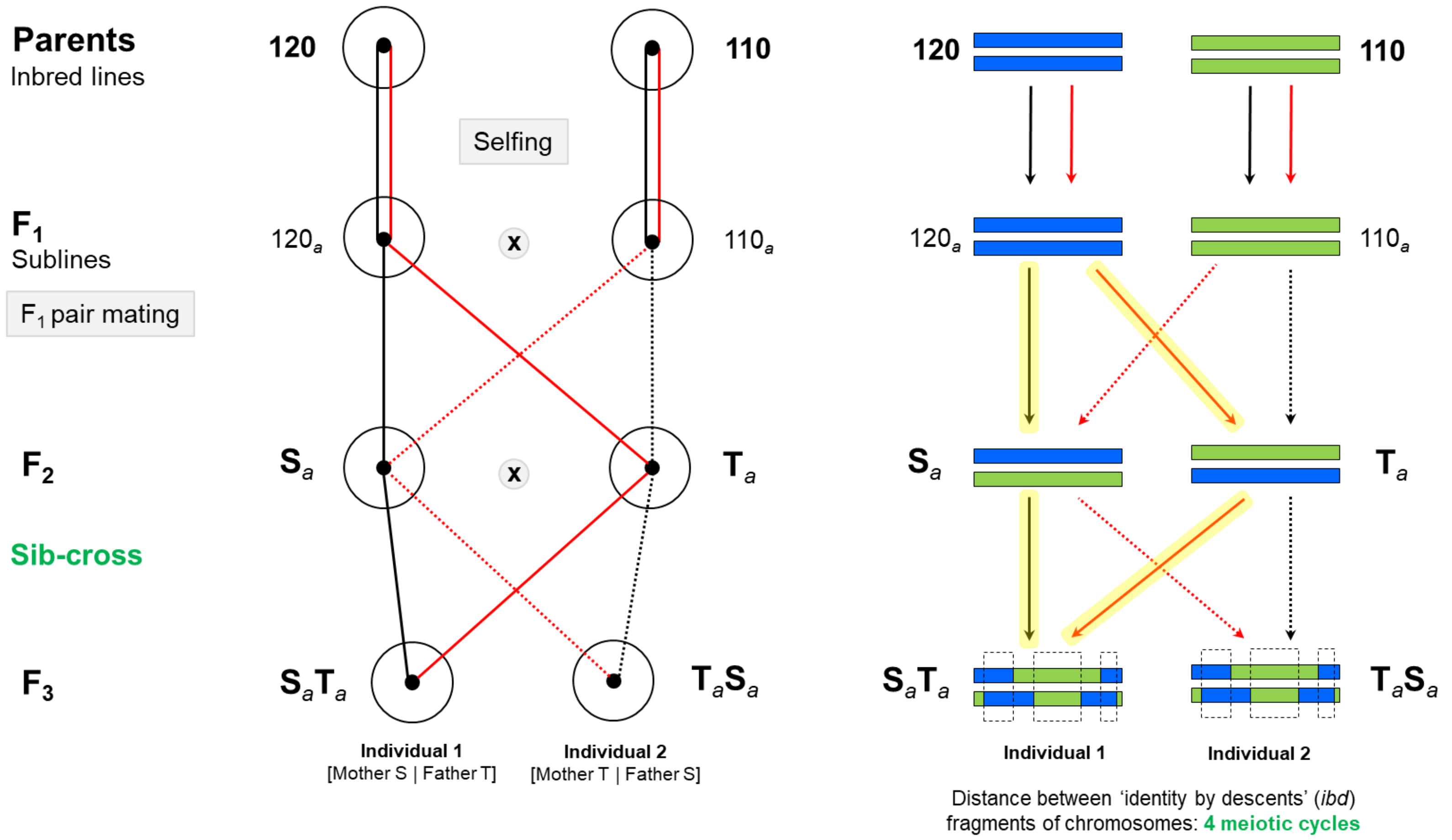

**Figure S5.** Pedigree of $F_3$ snails resulting from a one-way cousin-cross between two $F_2$ individuals of S origin belonging to different families (e.g., *a* and *b*), which are genetically identical to each other. Within *ibd* fragments the coalescence to common ancestral chromosome involves at least six meiotic cycles of which, on average, more than half are female meioses for "120" (blue) regions, and less than half are male meioses for "110" (green) regions (one possible pathway is highlighted in yellow).

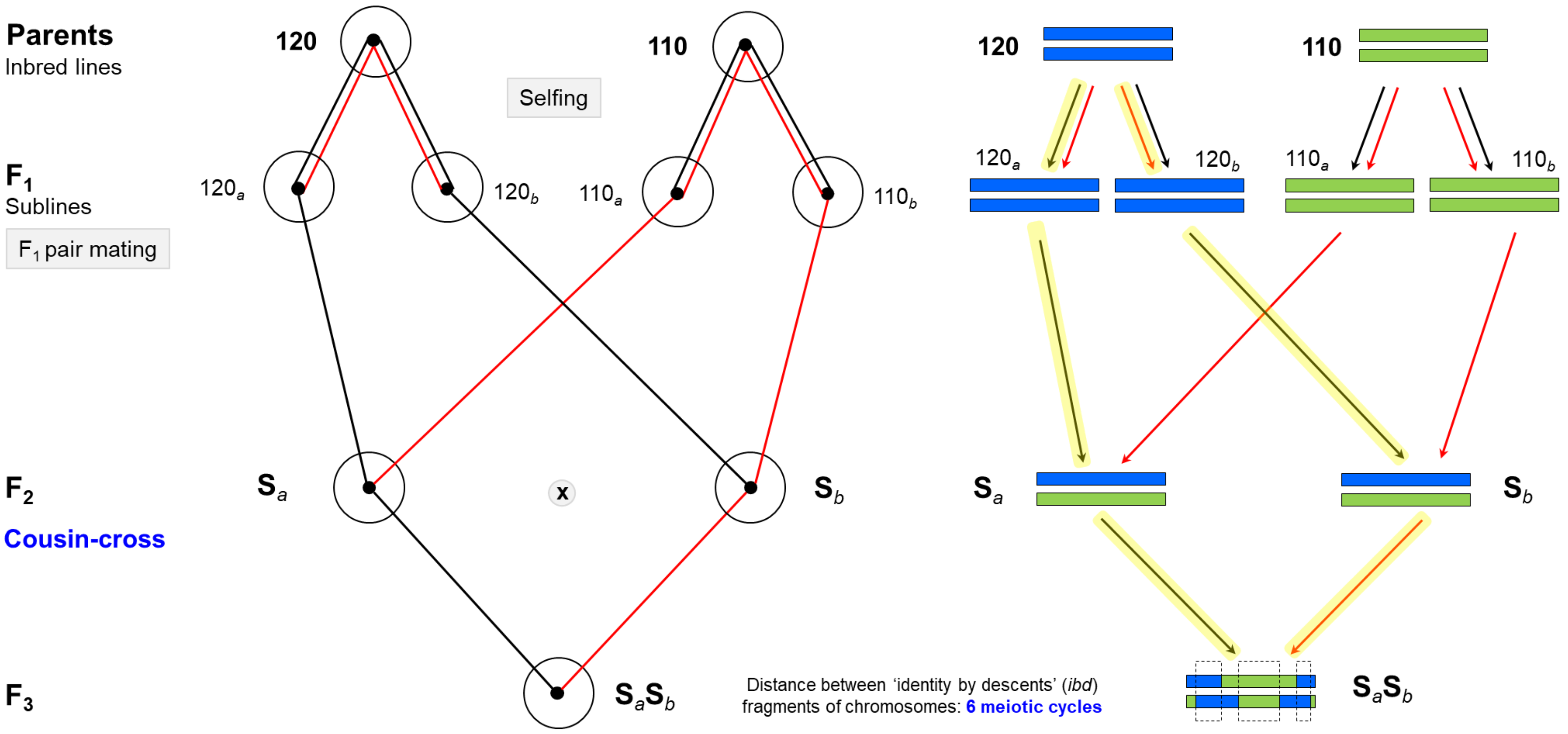

**Figure S6.** Pedigree of $F_3$ snails resulting from a two-way cousin-cross of $F_2$ individuals from different origins (S and T) and from different families (e.g., *a* and *b*). The distance between the *ibd* fragments to coalescence to the ancestral chromosomes (Parents) is at least six meiotic cycles; half of them are, on average, female meiosis both for "120" or "110" fragments (one possible pathway is highlighted in yellow).

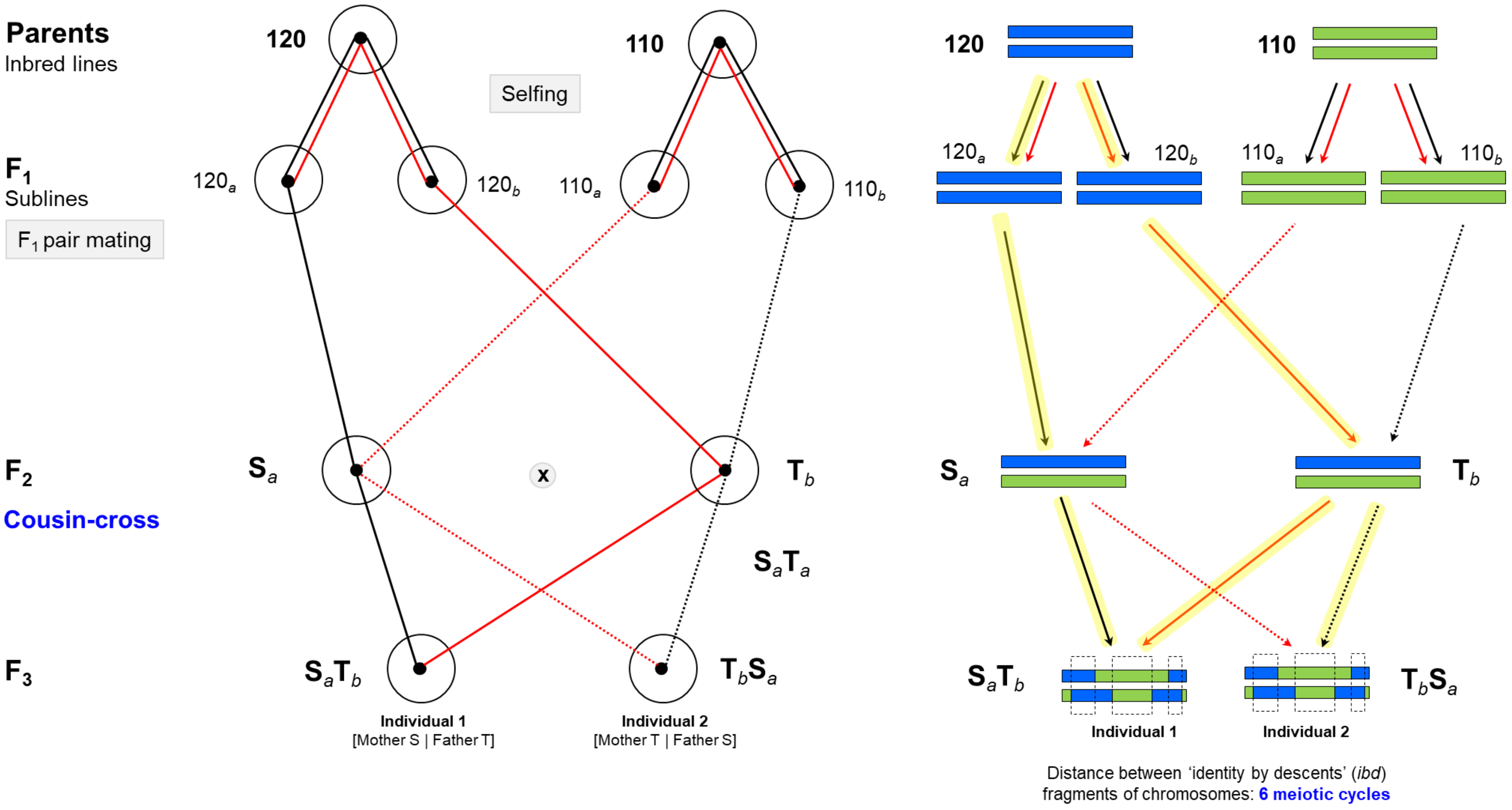

**Figure S7.** Simulations of the experiment under various parameter sets. The results are the means (± SD) of fitness ratios across 1000 simulations. The fitness ratio is computed as the ratio of the average fitness trait in offspring of sib- or cousin-cross to that of selfed offspring, and from cousin- to sib-cross. The parameters varied in the simulations are the contributions of semi lethal and small-effect mutations ($d_1$, $d_2$) to inbreeding depression in the base population, the genome length (*L* in Morgans), the dominance coefficient and effect of semi lethals mutations ($h_1$, $s_1$), and their equivalents for small-effect mutations ($h_2$, $s_2$). Each series of graphs from left to right represents the variation of one parameter, with all others held constant at reference values: $d_1$, $d_2$ = 0.3, 0.2; $L$ = 10; $h_1$ = 0.02; $h_1$ = 0.9; $h_2$ = 0. 2; $s_2$ = 0.05. The reference parameter set itself is represented as the third point in the first series but not repeated in the other series. The red and blue lines mark the observed fitness ratios for juvenile survival and body mass from the present experiment, respectively. The observed ratios for the probability of laying eggs at 50 days were Sibling/Selfing = 2.80, Cousin/Selfing = 4.30, and Cousin/Sibling = 1.54 (not shown in the figure, out of range).

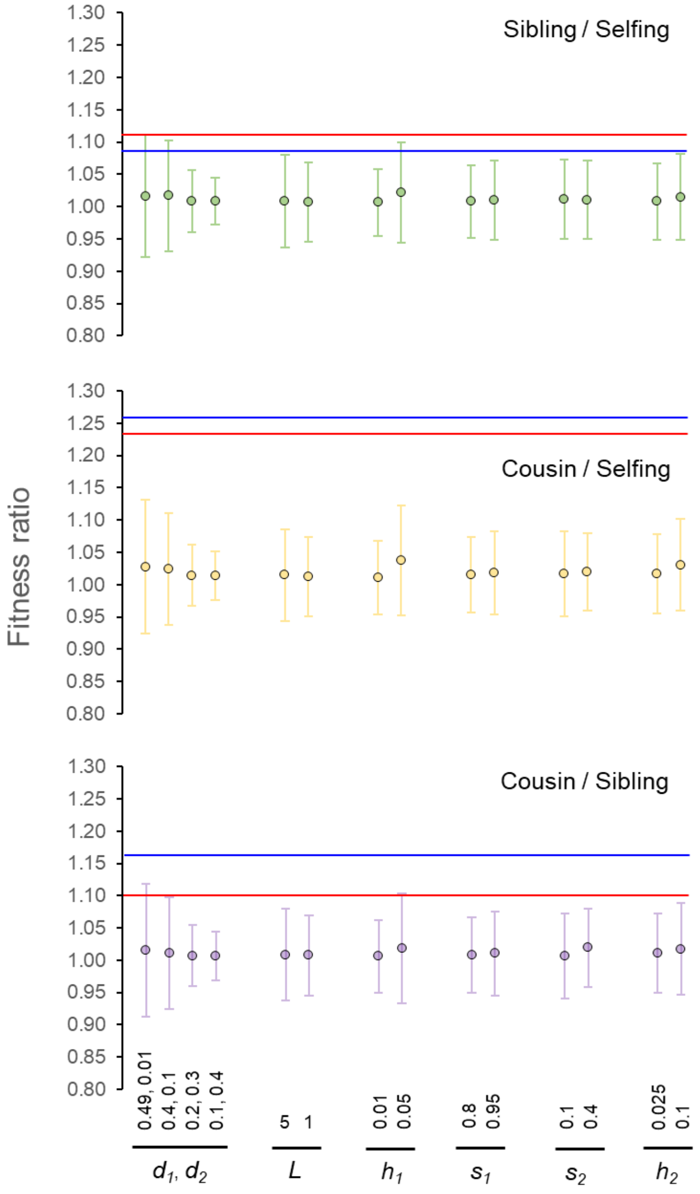

**Figure S8.** Distribution of fitness ratios from simulations with increased mutation rates. Plots show the fitness ratio values from 100 simulations of our experiment using the reference parameter set from Figure S7, with the mutation rates U1 and U2 progressively increased from one to 15 times the reference values. The fitness ratio is computed as the ratio of the average fitness trait in offspring of sibling-cross to selfed offspring, cousin-cross to selfed offspring, and cousin-cross to sibling-cross. Colored dots represent the mean fitness ratio values (± SD over simulations) for each mutation rate. The red and blue horizontal lines indicate the observed fitness ratios for juvenile survival and body mass from the present experiment. Observed fitness ratios for the probability of laying eggs at 50 days were Sibling/Selfing = 2.80, Cousin/Selfing = 4.30, and Cousin/Sibling = 1.54 (values out of the plotted range and not shown).

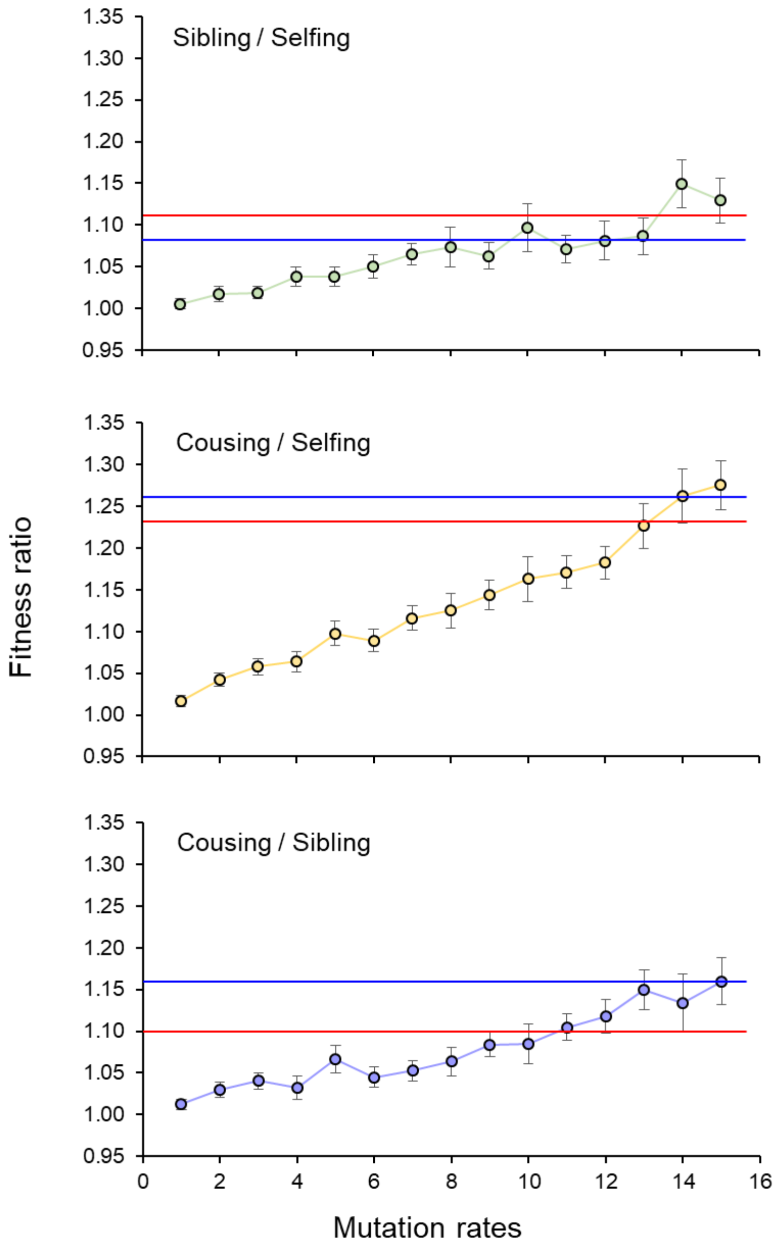
